# Modular striatal dynamics and reciprocal inhibition orchestrate skilled action sequencing

**DOI:** 10.1101/2025.06.27.662029

**Authors:** Konstantin Bakhurin, Bryan Lu, Hanchang Zheng, JongYup Park, Henry H. Yin

## Abstract

The basal ganglia, particularly the striatum, are critical for orchestrating skilled behavioral sequences, yet the precise mechanisms underlying this process remain unclear. Using high-resolution kinematic tracking and neural recordings in mice performing a water-reaching task, we found spatially distributed modules of striatal projection neurons whose activity corresponded to the generation of each element in the sequence—aiming, reaching, and drinking. They are activated sequentially and exhibit reciprocal inhibition, ensuring a strict serial order. Optogenetic activation of the direct pathway of the orofacial module promoted licking while suppressing reaching. Reaching could in turn suppress stimulation-evoked licking, revealing bidirectional inhibitory interactions. Our findings demonstrate that the striatum’s modular organization, coupled with lateral inhibition, sculpts the temporal progression of actions, providing a mechanistic framework for understanding how the basal ganglia coordinate complex behaviors.

## Introduction

Skilled behaviors involve combining multiple action elements into a precise sequence. The basal ganglia, particularly the striatum, are known to play a critical role in both action selection and sequencing^1–3^. The striatum receives major excitatory inputs from the cortex and thalamus^4,5^. In the sensorimotor regions, these inputs are organized somatotopically, with dorsolateral regions processing forelimb, trunk, and whisker-related behavior ^6–8^, and ventrolateral regions implicated in orofacial behavior^7,9,10^. The topographic organization suggests that actions recruiting different body parts activate distinct striatal regions^11,12^.

Research on striatal activity during goal-directed behavior reveals a strong link between behavioral kinematics, especially movement velocity, and basal ganglia function. Striatal projection neurons (SPNs) represent velocity, and stimulating these neurons can systematically modify movement kinematics ^13–16^. However, traditional studies of trained action sequences have often relied on limited behavioral measures, such as lever presses or beam breaks^17,18^, which fail to capture the detailed movement parameters and subtle transitions in behavior. While some studies have tracked movements in skilled tasks, these behaviors typically require extensive training, and the neural correlates of individual action components have not been thoroughly investigated^19,20^.

We recorded videos of mice performing forelimb reaching for water, while recording from different striatal regions using in vivo electrophysiology ^21^. Mice can easily learn this task, usually within hours, and in trained mice this behavior can be described as a simple sequence of three action components with clearly defined subgoals. Using this task, we discovered spatially segregated functional modules in the striatum, consistent with known topographic organization of the corticostriatal projections and striatal functional modules^12^. These functional modules are activated sequentially in time according to a strict serial order: aim, reach, drink. As neurons in a given module is suppressed while another module is active, our results suggest a reciprocal inhibition organization that allows the performance of one action component to suppress the expression of another, thus shaping the timing and ordering of action sequences.

## Results

### Effector alternation characterizes the performance of forelimb reaching

We trained freely moving mice to perform a reaching task for drops of water. The mouse has to reach through a narrow opening in a plexiglass arena^21^. Mice can quickly learn to reach water successfully within hours of training. This behavior is characterized by a sequence of head, forelimb, and orofacial movements (**Figure 1A**), each characterized by a distinct set of effector velocity profiles, which we called Aiming, Reaching, and Drinking. We quantified the kinematics of the head and the forelimb using DeepLabCut^22^. Aiming is characterized by an increased velocity of the head through the opening toward the water while the paw remains stationary (**Figure 1B**). During Reaching, the paw moves through the opening towards the drop of water. There was a concurrent reduction in head velocity during the increase in velocity of the paw. Lastly, Drinking involves bringing the paw holding the water to the mouth, as the mouse moves backward and starts licking its paw. The entire cycle lasts around 1250 ms (**Figure S1**).

**Figure 1.**
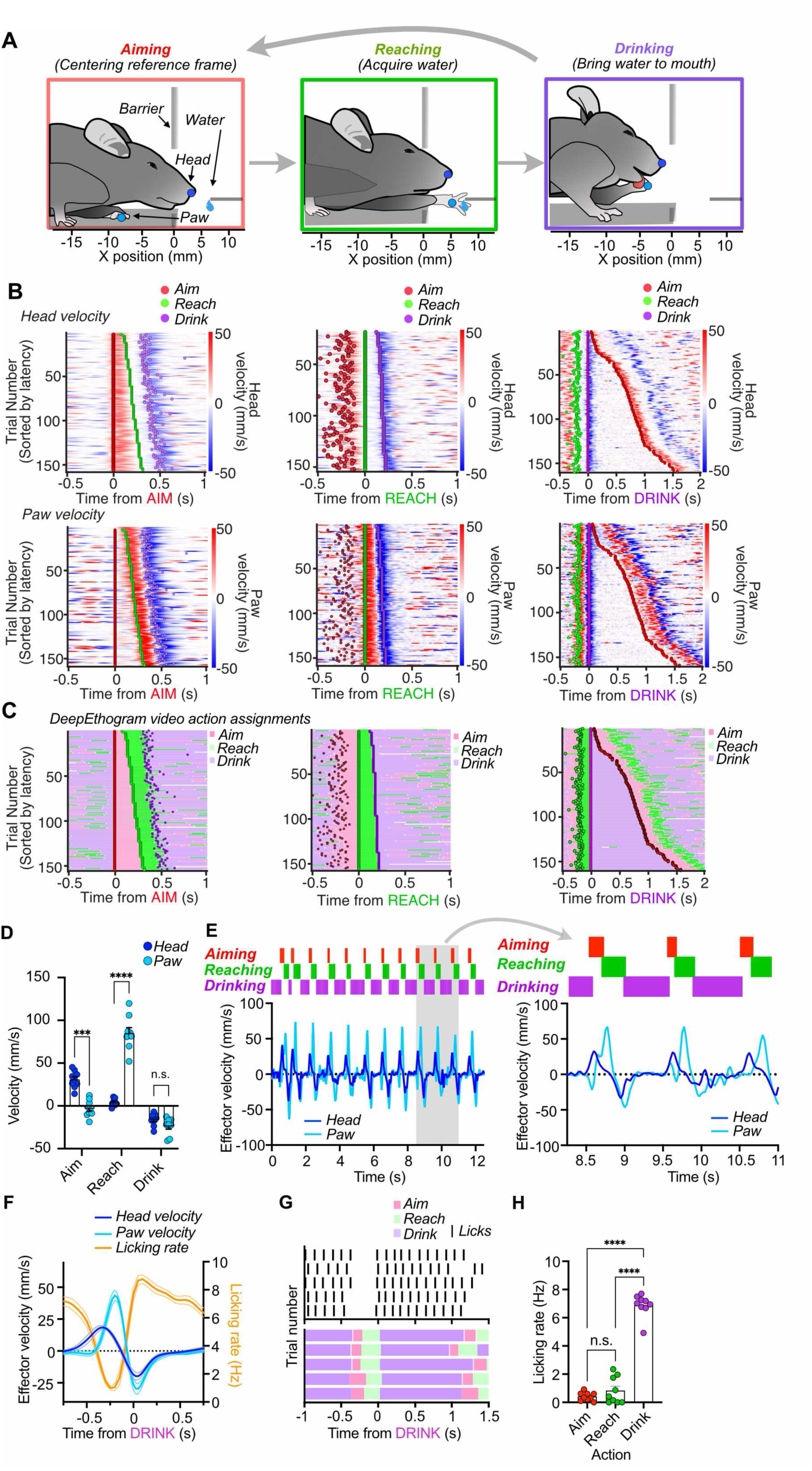
Reaching for water is a natural sequence. **A)** Freely moving mice generate a repeating sequence of posture transitions to perform a water reaching task. Aiming: movement of the head toward the target, Reaching: movement of the paw to grab water, Drinking: water is brought to the mouth by the paw and is consumed. **B**) Heatmaps for head (*Top*) and paw (*Bottom*) velocity aligned to Aim, Reach, and Drink actions for a single session. Trials are sorted by time to the next action. **C**) Diagrams showing Deepethogram labels for Aim, Reach, and Drink corresponding to the same data as in *B*. **D**) There is an alternation in velocity of the head and paw during the transition to Aim and to Reach. (Two-way RM ANOVA, Main effects of Action (F(2,16) = 125.5, p < 0.0001) and Effector (F(1,8) = 60.7, p < 0.0001),significant interaction (F(2,16) = 98.46, p < 0.0001). Post-hoc tests revealed that Head velocity was greater during Aiming (p = 0.0001), Paw velocity was greater during Reaching (p < 0.0001), and no difference during Drinking (p = 0.5). **E**) *Left*: Ethogram showing distinct action identification from video, with each frame being assigned to one action. *Right*: Same data but zoomed in to the time period depicted in the gray rectangle at left. Traces below show the alternating velocity profiles of the effectors in Aiming and Reaching, the head and the paw. **F**) Average licking rate overlayed with Head and Paw velocity aligned to the onset of Drinking (n = 9 mice). Licking is reduced as the Aim and Reach are initiated. **G**) Representative raster of individual licks aligned to 5 Drinking actions. **H**) Mean licking rate during the three actions observed during water reaching behavior. A one-way repeated measures ANOVA found a significant effect of action component on licking rate (F(1.98,15.91) = 216, p < 0.0001). Post-hoc tests found that licking rates were much higher during Drinking compared to Aiming (p < 0.0001) and Reaching (p < 0.0001).

We used a deep neural network pipeline (DeepEthogram) to classify each video frame as belonging to either Aiming, Reaching, or Drinking (**Figure 1C**)^23,24^. Aligning the kinematics to Aiming, Reaching, and Drinking labels shows that the movements of the head and paw operate in a serial process, alternating velocity profiles over time (**Figure 1D & 1E**). Head and paw movements were not simultaneous as the paw lagged the head by approximately 150 ms (**Figure S1**). Lastly, after grabbing the water, licking increased as the head and paw moved backward to drink. Licking was again suppressed as mice began the next Aim and would remain inactivated until initiating Drinking again (**Figures 1F & 1G**). Licking rates were low during Aiming and Reaching but high during Drinking (**Figure 1H**). Together, reaching for water requires a coordinated sequence of actions, each associated with a distinct set of transient effector movements (**Video S1**).

### Distributed modular representations of actions in the sensorimotor striatum

We mapped the functional roles of projection populations along the dorsoventral axis of the sensorimotor striatum during reaching. We implanted chronic, drivable micro-electrode arrays above either the DLS or the VLS in D1-Cre and A2A-Cre mice (**Figure 2A**). We isolated putative SPNs (n = 572 from VLS mice, 5 mice, 1075 from DLS, 7 mice) by using clustering of waveform shapes projected onto PCA space (**Figure S2**). SPNs displayed typical low firing rates and wide waveforms. DLS SPNs and VLS SPNs showed reciprocal patterns of activation during reaching (**Figure 2B**). DLS firing rates increased prior to Reaching, while the VLS firing rates reduced. When mice transitioned to Drinking, VLS activity increased while DLS activity was reduced (**Figure 2C**). These results reveal that these striatal domains are differentially active during different phases of the reaching sequence.

**Figure 2.**
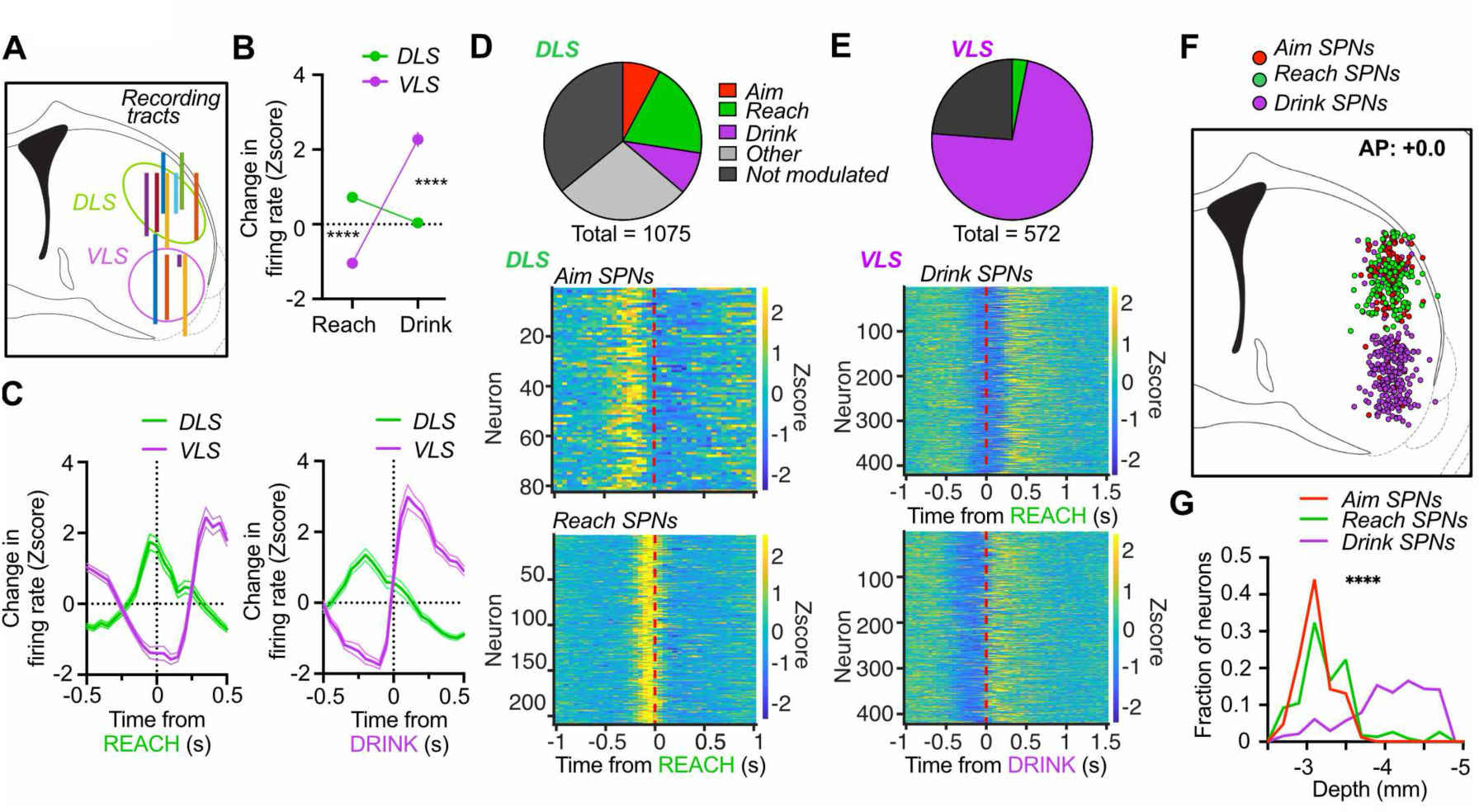
Opponent activation of dorsal and ventral striatal neuron populations during reaching. **A**) Diagram showing drivable electrode tracts in the DLS (n = 7 mice) and VLS (n = 5 mice). **B**) Opponent activation by DLS and VLS SPNs during Reaching and Drinking (Two-way ANOVA, Main effect of Action (F(1,1504) = 108.5. Significant interaction between Region and Action (F(1,1504) = 251.7). Post-hoc tests found that DLS activity was greater during Reaching (p < 0.0001) and VLS activity was greater during Drinking (p < 0.0001). **C**) Mean firing rate for DLS and VLS SPNs aligned to Reaching (*Left*) and Drinking (*Right*). **D**) Proportion of all SPNs that belonged to Aiming, Reaching, and Drinking groups in the DLS. Heatmaps show the population activity of A-SPNs (*Middle*, n = 84) and R-SPNs (*Bottom*, n = 211) in DLS aligned to Reach onset. **E**) Proportion of all SPNs that belonged to Reaching and Drinking groups. Heatmaps show population activity of D-SPNs aligned to Reach onset (*Middle*) and Drink onset (B*ottom*, n = 418 SPNs). **F**) Coronal view of the striatum showing the locations all A-, R-, and D-SPNs. **G**) Distributions of depths of all A-, R-, and D-SPNs (Significant one-way ANOVA (p < 0.0001). D-SPNs are deeper than A-(p < 0.0001) and R-SPNs (p < 0.0001). R-SPNs are deeper than A-SPNs (p = 0.018)).

We made further classifications of DLS and VLS SPN populations by aligning their activity to Reaching and performing hierarchical clustering on the population activity (**Figure S2**). We identified three task-relevant populations. The DLS contained large populations of Aiming SPNs (A-SPNs) and Reaching SPNs (R-SPNs) populations. These were active prior to Reaching and had phasic activations of different latencies (**Figure 2D**). VLS SPNs largely belonged to a Drinking-related population (D-SPNs) whose activity was suppressed during the reach but showed sustained firing activity during Drinking (**Figure 2E**). We found that Aiming and Reaching neurons were most prominently found in the DLS, whereas Drinking neurons were more prevalent in the VLS (**Figure 2F & G**).

### Alternation and suppression of SPN modular activity during transition initiation

We analyzed the relationships between A-SPNs and R-SPNs and kinematics in the task. A-SPNs fired as mice increased their head velocity to aim at the target (**Figure 3A**). Their transient pattern of activity resulted in an average firing rate profile that matched the velocity of the head (**Figure 3B**). The neurons were tuned to the velocity of the head toward the target, but not during movements of the head away from the target, reflecting a rectified tuning property. A-SPN activity led both the head and the paw velocity, indicating that these neurons were active prior to head movements (**Figure 3C**). A-SPNs showed increases in firing rate prior to Aiming onset but showed reductions in firing rate prior to Reaching and during Drinking (**Figure 3D**).

**Figure 3.**
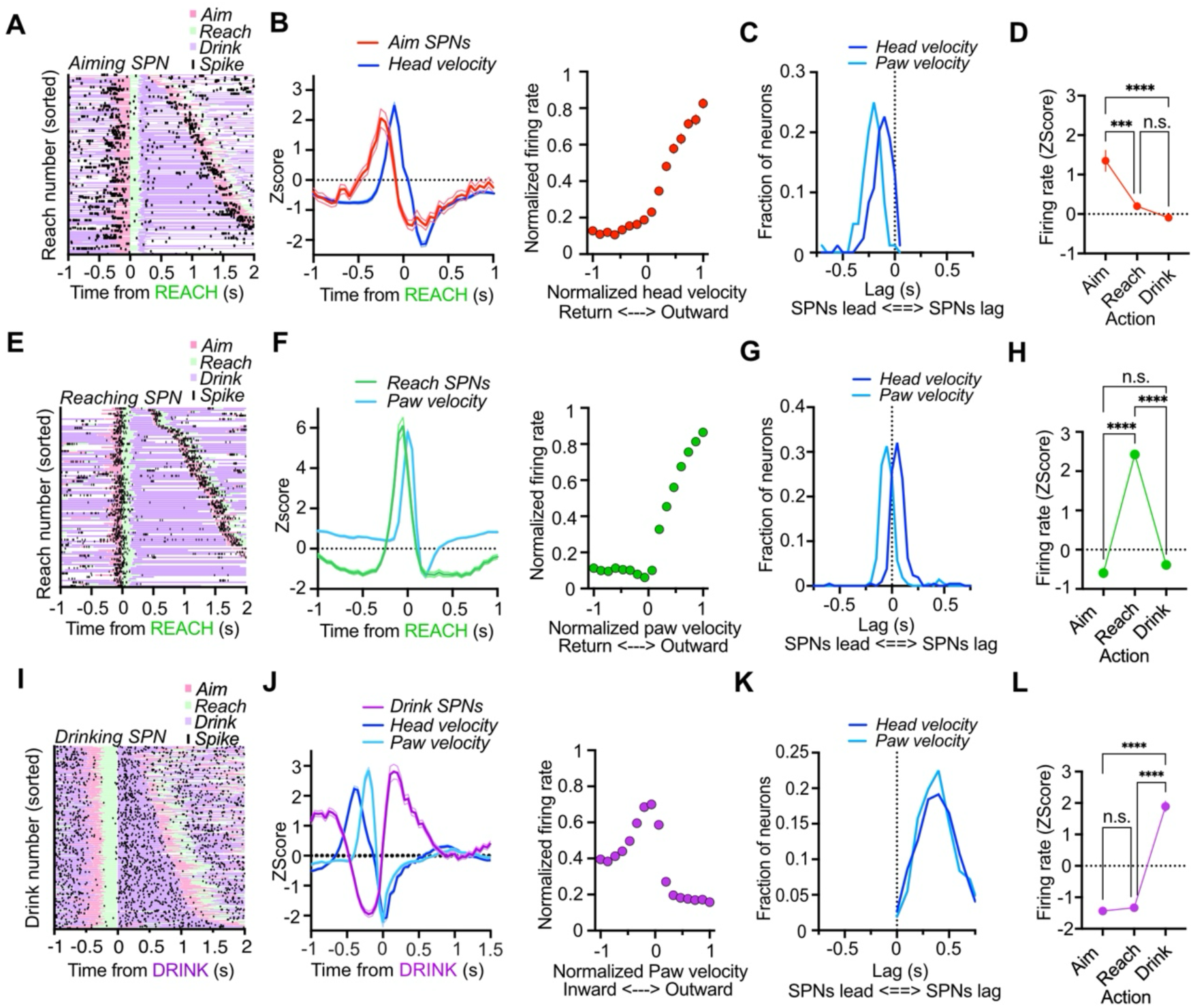
Alternating activation and suppression of action-specific striatal modules during the reaching sequence. **A**) Example raster plot of an A-SPN showing bursting prior to onset of Aim. **B**) Population average of A-SPNs overlayed with head velocity (n = 84 SPNs from 7 mice). *Right*: Average tuning curve for A-SPN population firing vs head velocity (n = 84 SPNs). **C**) A-SPNs lead both head and paw velocity. **D**) Average firing rates for A-SPNs aligned to Aim, Reach, and Drink (One-way repeated measures ANOVA, F(1.31, 108.8) = 21.11, p < 0.0001). A-SPNs fired more during Aiming than Reaching (p = 0.0004) and Drinking (p < 0.0001). **E**) Example raster plot of an R-SPN showing bursting prior to onset of Reach. **F**) Population average of R-SPNs overlayed with paw velocity (n = 211 SPNs from 7 mice). *Right*: R-SPN population firing vs paw velocity (n = 211 SPNs). **G**) R-SPNs lead paw velocity and lag nose velocity. **H**) Average firing rates for R-SPNs aligned to Aim, Reach, and Drink (One-way repeated measures ANOVA, F(1.3, 274) = 497.7, p < 0.0001). R-SPNs fired more during Reaching than Aiming (p < 0.0001) and Drinking (p < 0.0001). **I**) Example raster plot of a D-SPN in the VLS, aligned to Drink onset. Trials are sorted by latency to Drink offset. **J**) *Left*: Mean activity of D-SPNs aligned to Drink plotted together with Head and Paw velocity. D-SPNs reduce their activity during Reaching. *Right*: D-SPN population activity as a function of paw velocity. **K**) Distribution of cross-correlation lags relative to paw velocity. Absolute value is shown. **L**) D-SPNs are suppressed during Aiming and Reaching but activate during Drinking (One-way RM ANOVA, F(1.52, 633.6) = 281.2, p < 0.0001). Post-hoc tests: Firing during Drink is higher than Reach (p < 0.0001) and Aim (p < 0.0001). Reach is also higher than Aim (p < 0.0001).

R-SPNs fired when mice generated movements of the arm to obtain the water (**Figure 3E**). These neurons’ firing rate matched the velocity profile of paw movement (**Figure 3F**). R-SPN neuron activity was correlated with the velocity of the arm as it approached the target, but not when the arm was returning into the arena, also displaying rectified tuning (**Figure 3F**). R-SPN activity increased just prior to paw velocity but lagged behind increases in head velocity (**Figure 3G**). R-SPNs showed higher firing activity prior to Reaching but were reduced in firing below baseline prior to Aiming and during Drinking (**Figure 3H**).

In contrast to the A– and R-SPN modules, D-SPNs are active during Drinking (**Figure 3I**). D-SPNs show an antiphase relationship to head paw velocity, showing an increase when aligned to Drinking but a suppression during Aiming and Reaching. D-SPN firing rates were therefore greatest at the lowest paw velocities while mice were consuming the water (**Figure 3J**). Activity in D-SPNs lags significantly behind reaching (**Figure 3K**). Drinking neuron activity was reduced when mice initiated Aiming and Reaching, but during Drinking, D-SPNs increase their activity (**Figure 3L**). Together, we found discrete populations of SPNs in the DLS that were associated with transitions of different goal-directed components of the reaching sequence. Each population activated during their respective action but was suppressed when a different action was being performed.

### Dynamic antagonism between modules sculpts sequence progression

Because Deepethogram labels reflected the transition into a specific action, they could be used to represent and directly compare the temporal relationships between actions during the reaching behavior, even between animals. Thus. when mice were Reaching, there was a low probability of observing Drinking labels and vice versa (**Figure 4A & B**). These labels allowed us to align neural activity from the VLS and the DLS to the onset of Reaching, confirming that these populations alternate in their activity (**Figure 4C**).

**Figure 4.**
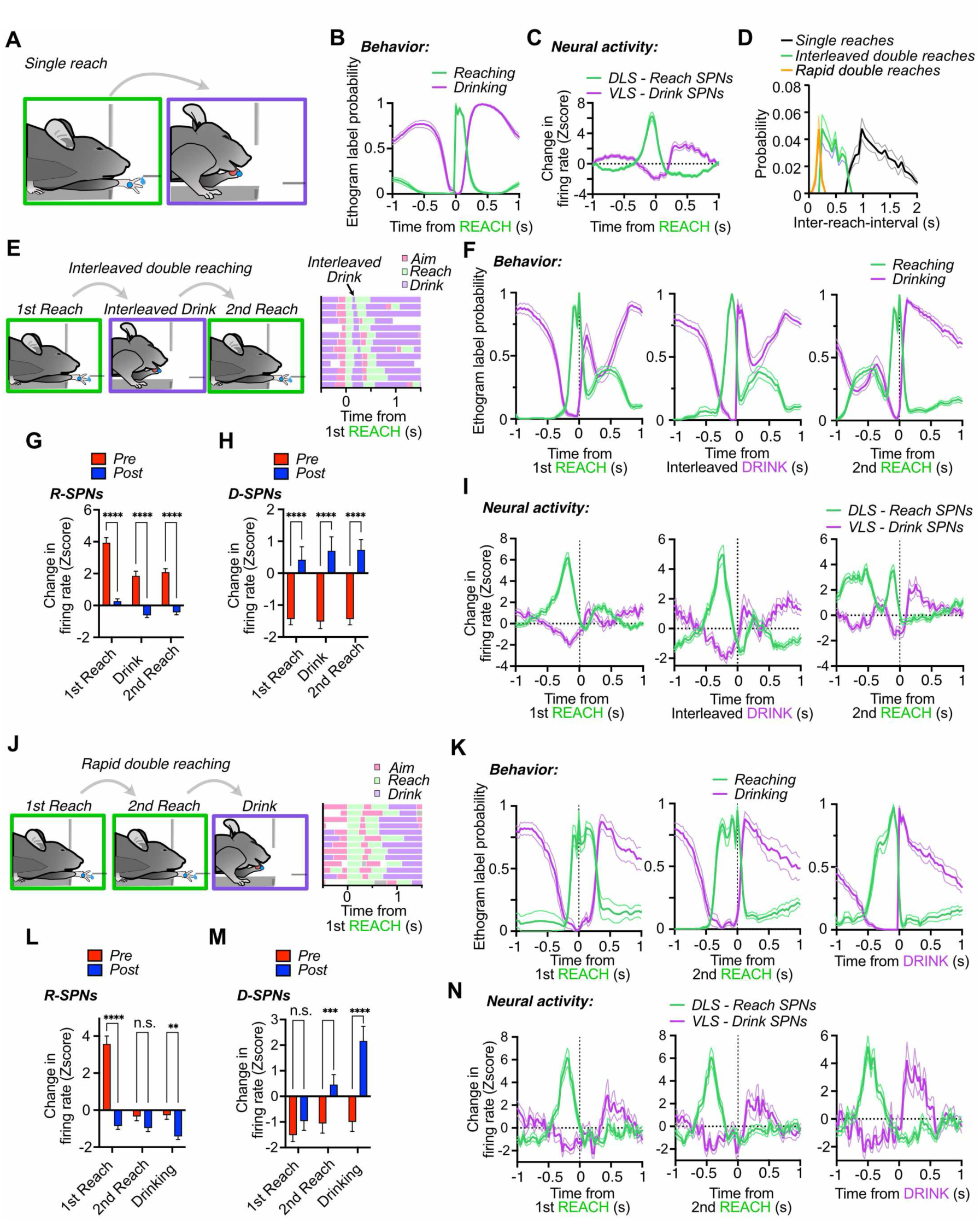
Antagonism between Reaching and Drinking systems sculpts sequence progression. **A**) Diagram showing the normal sequence of reach and drink transitions. **B**) Mean Reach and Drink Deepethogram label probability aligned to reach onset (n = 11 mice). **C**) Mean Reach (n = 121) and Drink (n = 81) SPN firing rates aligned to single reach onset (n = 11 mice). **D**) Distribution of inter-reach-intervals, showing separation of Double and Single reaches. Interleaved-double reaches are those with inter-reach-intervals between 250 and 750 ms. Rapid double reaches were those with inter-reach-intervals less than 250 ms. **E**) Diagram showing Interleaved-Double reaches in which mice briefly lick the paw between two reaches in quick succession. *<u>Right</u>*: example Deepethogram aligned to first reach onset showing the brief drink action. **F**) Deepethogram label probability aligned to the first reach, the interleaved drink, and the second reach of Interleaved double reach events (n = 11 mice). Note reciprocal alternation of action labels. **G**) Changes in firing rates of Reaching (n = 121) SPNs during Interleaved-double trials. Mean rates were calculated prior to (pre: –250 to 0 ms) and after (post: 0 to 250ms) the first reach, the interleaved drink, and the second reach of Interleaved-double trials. Two-way RM ANOVA, significant main effects of Action (F(2,240) = 43.27, p < 0.0001) and Analysis interval (F(1,120) = 154, p < 0.0001), and a significant interaction (F(2,240) = 10.51, p < 0.0001). Post-hoc tests showed a significant reductions in R-SPN firing rate immediately after the 1^st^ reach (p < 0.0001), the interleaved drink (p < 0.0001), and the 2^nd^ reach (p < 0.0001). **H**) Changes in firing rates of Drinking (n = 81) SPNs during Interleaved-double trials. Mean rates were calculated prior (pre: –250 to 0 ms) and after (post: 0 to 250ms) the first reach, the interleaved drink, and the second reach of Interleaved-double trials. Two-way RM ANOVA, significant main effect of analysis interval (F(1,80) = 41.62, p < 0.0001), but no significant interaction (F(2,160) = 0.6, p = 0.54). D-SPNs showed significant increases in firing after each action (p < 0.0001). **I**) Firing rates of Reaching and Drinking SPNs aligned to the first reach, the interleaved drink, and the second reach of Interleaved-double trials. Note alternating patterns of activity that corresponds to the action components in *F.* **J**) Diagram showing Rapid double reaches that are not interrupted by a drink event. *Right:* example Deepethogram for Rapid double reaches. **K**) Mean Deepethogram labels for Reaching and Drinking aligned to the first reach, second reach, and drinking action of rapid double reach trials (n = 11 mice). **L**) Changes in firing rates for Reach (n = 121) SPNs during Rapid-double reaching. Mean rates were calculated prior to (pre: –250 to 0 ms) and after (post: 0 to 250ms) the first reach, the second reach, and the final drink action of these trials. Two-way RM ANOVA, significant main effects of Analysis interval (F(1,120) = 93.37, p < 0.0001) and Action (F(2,240) = 70.94, p < 0.0001) and a significant interaction (F(2,240) = 40.54, p < 0.0001). Post-hoc tests showed that firing rates were significantly increased before the first reach (p < 0.0001) and suppressed after drinking (p = 0.0013). **M)** Changes in firing rates for Drinking (n = 81) SPNs during Rapid-double reaching. Mean rates were calculated prior to (pre: –250 to 0 ms) and after (post: 0 to 250ms) the first reach, the second reach, and the final drink action of these trials. Two-way RM ANOVA, significant main effects of Interval (F(1,80) = 33.39, p < 0.0001) and Action (F(2,160) = 23.03, p < 0.0001). There was a significant interaction (F(2,160) = 11.97, p < 0.0001). Post –hoc tests found a significant increase in firing after the second reach (p = 0.0003) and after Drinking (p < 0.0001). **N**) Activity of Reaching and Drinking SPNs aligned to the first reach, second reach, and drinking action of Rapid-double trials. Note reduced activity of both modules during the generation of the second reach.

Although mice usually produced successful single reaches that were followed by Drinking of the water, mice occasionally produced repeated reach events in quick succession (**Figure 4D, Video S2**), represented by low inter-reach-intervals. Double reaches contained two reach events within 750ms of one another, whereas single reaches were separated by a longer drinking bout. In most cases, the multiple reaches in a double reach trial were separated by a brief, interleaved drinking event, possibly for mice to check if they had obtained any water (**Figure 4E & Figure S3**). These types of trials could be isolated by taking reaches that occurred within 250 and 750ms of each other. Rapid Double reaches occurred within 250ms of each other and were not interrupted by any Drinking, reflecting a quick succession of two reach events close in time that delayed Drinking (**Figures 4D, 4J, Figure S3**). Both types of double reach trials allowed us to explore the dynamic interaction between Reaching and Drinking modules in two different sequences.

During Interleaved-double trials, reaches were interrupted by a brief Drinking action before the next reach was repeated (**Figure 4E**). The latency to initiate the interleaved Drinking action was not different from latency to initiate Drinking during single reaching, but the duration of these interleaved drink actions was shorter (**Figure S3**). Paw velocity during Interleaved-double trials exhibited two peaks when aligned to each of the two reaches (**Figure S3**). By aligning Deepethogram labels to the first reach, Interleaved drinking, and the second reach, we could observe an alternating pattern of Reaching and Drinking actions during these trials (**Figure 4F**). R-SPN activity increased prior to each action in the sequence and was reduced immediately after (**Figure 4G**). In contrast, D-SPN activity was suppressed prior to the actions, but increased after (**Figure 4H**). Neural activity in R– and D-SPNs thus reflected an alternating pattern that paralleled the dynamic relationships between Reaching and Drinking (**Figure 4I**).

During Rapid double reaching, mice produced two reaches without any Drinking event interrupting them (**Figure 4J**). This was reflected by two peaks in paw velocity occurring close together (**Figure S4**). Reaching action duration was extended to accommodate the multiple reaches and the latency to initiate Drinking was delayed (**Figure 4K & Figure S3**). R-SPN activity prior to the first reach was increased and was suppressed for the remainder of the sequence (**Figure 4L**). D-SPN activity was suppressed until after completion of the second reach (**Figure 4M**). We therefore found that R-SPN activity can initiate the production of multiple repeated movements without firing multiple times, and that D-SPN activity is delayed until after the reaching system is deactivated (**Figure 4N**). While Reaching action is being produced, Drinking is delayed. However, the brief activation of the Drinking module during Interleaved trials necessitates an increase in Reaching module activity prior to the second reach. Together these results reveal how action modules are flexibly coordinated and recruited to accommodate changes in the basic sequence, indicating the striatum actively determines sequence order and progression.

### VLS direct pathway promotes licking while suppressing reaching

Because the VLS contained a segregated neural population that was active during Drinking, we targeted the direct pathway (D1+ neurons) in this region to selectively activate the dedicated Drinking module (**Figure 5A & Figure S4**). We used the infrared beam break by the nose that occurs when mice perform the Aiming action to immediately trigger 30 Hz laser stimulation (**Figure 5B**). VLS stimulation at 30 Hz resulted in licking behavior outward toward the spout (**Figure 5C, Video S3**). Licking normally was delayed until after reaching, but stimulation reduced the latency to initiate licking (**Figure 5D & E**). We called these types of trials “Licking Wins” trials. During stimulation on these trials, head and paw were both immobile (**Figure 5F & Figure S4**). Whereas normally mice perform a reach and then withdraw the head after retrieving the water, direct pathway activation suppressed the head-retraction movement, instead maintaining their head in the reaching port (**Figure S4**). At the same time, paw velocity was reduced and reaching by the paw that occurs after Aiming was not observed (**Figure 5G**).

**Figure 5.**
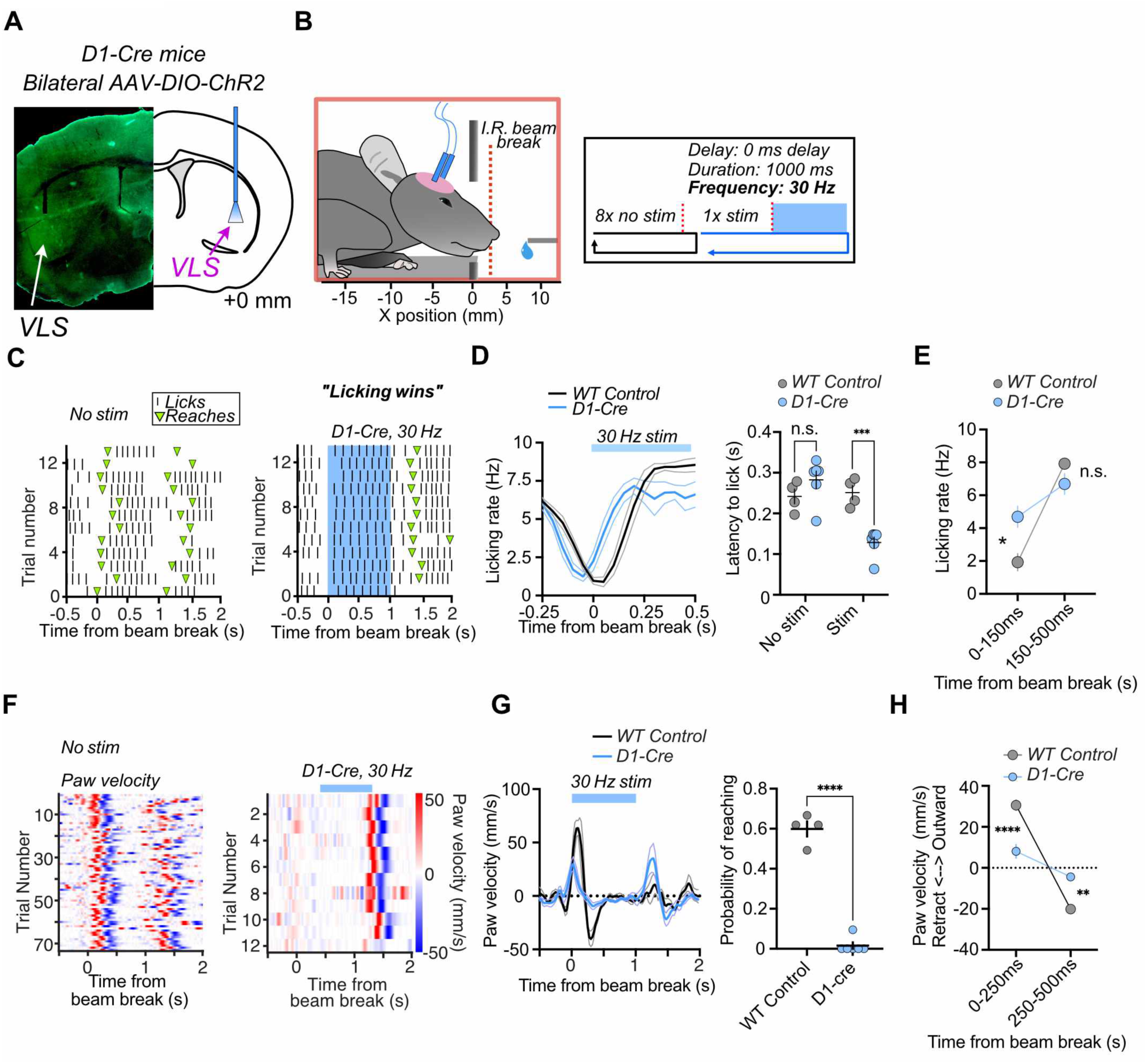
– Direct pathway activation in the VLS promotes licking while suppressing reaching. **A)** Expression of GFP in the VLS in a D1-cre mouse. **B**) Design. 30Hz simulation occurred for 1 second every 9^th^ beam break with the nose to deliver bilateral light stimulation to the VLS during Aiming. **C**) *Left*: Lick raster plot of a representative D1-cre mouse not receiving stimulation. *Right*: Raster plot showing licking evoked from VLS D1 pathway activation. Reaches (Green markers) are delayed until after stimulation. **D**) *Left*: Average licking rate time series during trials with light delivery to the VLS in D1-cre (n = 6) and WT mice (n = 4). *Right*: Average lick onset latency is shorter for trials with VLS D1 activation. Two-way ANOVA, significant main effect of Stimulation on lick latency (F(1,8) = 89.64, p < 0.0001) and a significant interaction between Stimulation and Group (F(1,8) = 114.3, p < 0.0001). Post-hoc tests showed that stimulation of D1 neurons reduced lick latency (p = 0.0006). **E**) Licking rate is increased 150ms after stimulation onset in D1-cre mice. A two-way ANOVA showed a significant main effect of Time Window (F(1,8) = 79.3, p < 0.0001) but not Group (F(1,8) = 1.1, p = 0.32) on licking rate. There was a significant interaction between Time Window and Group (F(1,8) = 19.8, p = 0.0021). Post-hoc tests showed that licking rate was higher in D1-cre mice during the first 150 ms after stimulation onset (p = 0.0104) but not in the subsequent time window (p = 0.31). **F)** *Left*: Representative heatmap showing normal paw velocity during reaching. *Right*: Representative heatmap showing the reduction of paw velocity during VLS direct pathway activation. **G**) *Left*: Average paw velocity time series of all subjects showing that direct pathway stimulation reduces paw movement (n = 6 D1-cre and 4 WT Control mice). *Right:* Reaching probability during the first second after stimulation was significantly reduced during direct pathway activation (unpaired t-test, p < 0.0001). **H**) VLS direct pathway activation suppresses paw movement during reaching. Two-way ANOVA, significant main effect of Time point (F(1,8) = 146.1, p < 0.0001) on paw velocity and a significant interaction between Time point and Experimental group (F(1,8) = 53.53, p < 0.0001). Post-hoc tests found significant reductions in paw velocity by stimulation of the VLS in D1-cre mice during the first (p < 0.0001) and second (p = 0.001) 250ms epochs of the reach.

The outward and the retraction velocity components of the reach were reduced (**Figure 5H**). Altogether, stimulation produced a continuous licking bout that suppressed the ability for mice to use their paw.

### Reaching suppresses direct-pathway induced licking behavior

We examined the effect of reducing laser frequency on the ability for VLS direct pathway to suppress reaching (**Figure 6A**). Reducing the stimulation frequency reduced the frequency of “Licking Wins” trials (**Figure 6B**). We found that reducing stimulation frequency revealed a greater likelihood of trials where mice could produce a successful reach (“Reaching Wins” trials, **Figure 6C**). For all stimulation frequencies, we found that paw velocity was intact during Reaching Wins trials but was suppressed during Licking Wins trials (**Figure 6D**). Licking latency was significantly reduced during Lick Wins trials but was normal for Reaching Wins trials (**Figure S5**). In contrast, reach latency was increased during Lick Wins trials, but normal for Reach Wins trials (**Figure S5**). Overall, stimulation frequency did not significantly influence these changes in movement latencies during Licking Wins and Reaching Wins trials but only influenced the relative likelihoods of observing these trial types.

**Figure 6.**
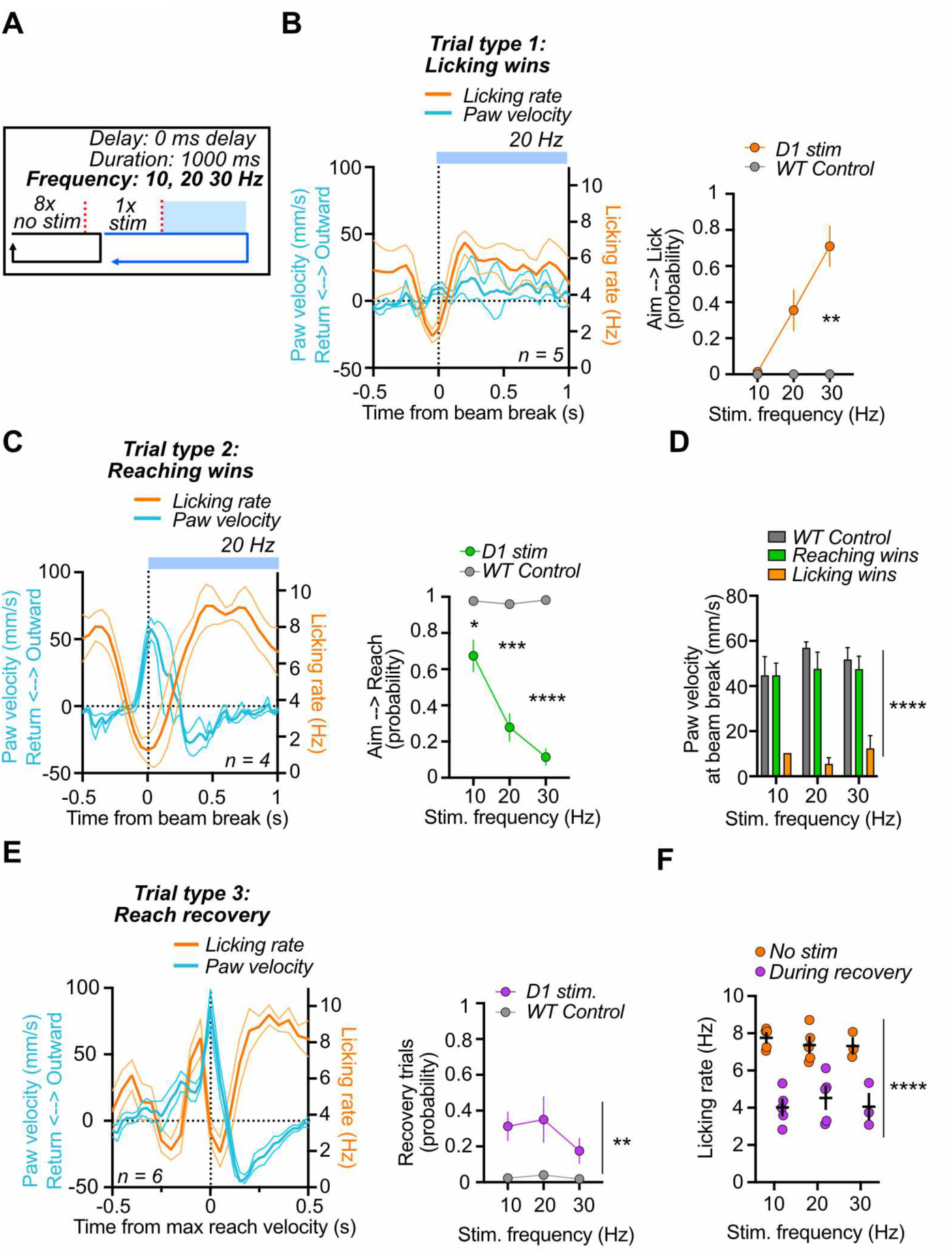
– Reaching during direct pathway stimulation suppresses light-evoked licking. **A**) Design. 10, 20, or 30Hz simulation occurred for 1 second every 9^th^ beam break with the nose to deliver bilateral light stimulation to the VLS during Aiming. **B**) Licking Wins trials. Reaching is suppressed if licking is initiated by stimulation. *Right*: The probability of observing Licking-wins trials increased with increasing stimulation frequency. Two-way ANOVA, main effects of Group (F(1,8) = 19.2, p = 0.0023), Frequency (F(1.95, 15.64) = 12.18, p = 0.0007) and a significant interaction between Group and Trial Type (F(2,16) = 12.18, p = 0.0006). Post-hoc tests for multiple comparisons found a significant increase in the probability of observing Licking wins trials during 30 Hz (p = 0.0044) stimulation in D1-cre mice. **C**) Reaching Wins trials. Licks are delayed if Reaching is produced during stimulation. *Right*: The probability of observing Reaching-wins trials was reduced with increasing stimulation frequency. Two-way ANOVA, main effects of Group (F(1,8) = 122, p < 0.0001), Frequency (F(1.1, 8.9) = 10.74, p = 0.0087) and a significant interaction between Group and Trial Type (F(2,16) = 10.7, p = 0.0011). Post-hoc tests for multiple comparisons found significant reductions in the probability of observing Reaching wins trials during 10 Hz (p = 0.048), 20 Hz (p = 0.0004), and 30 Hz (p < 0.0001) stimulation in D1-cre mice. **D**) The velocity at beam break time is significantly lower if stimulation evoked licking. A two-way ANOVA found a significant effect of Trial type on paw velocity at beam break (F(2,29) = 27.27, p < 0.0001) **E**) Licking rate and paw velocity overlayed and aligned to the peak of the Recovery reach velocity. Licking is reduced once a high paw velocity is performed. The probability of observing Recovery trials was not affected by frequency (Two-way ANOVA, main effect of Trial type (F(1,8) = 14.2), p = 0.0055). **F**) Licking rates are reduced during Recovery reaches (Two-way ANOVA, main effect of Trial type F(1,10) = 119, p < 0.0001).

On some trials, VLS direct pathway stimulation initially produced licking that was suppressed by the production of a Recovery reach during stimulation (**Figure 6E, Video S4**). There was no effect by stimulation frequency on the rate at which we observed these types of trials (**Figure 6E**). We detected the peak Reaching velocity during these trials when mice could overcome stimulation-evoked licking. By aligning to these high velocity movements, we found that stimulation-evoked licking was inversely related to the peak velocity (**Figure 6F**). There was a reduction in licking latency and an increase in reaching latency (**Figure S5**). Thus mice could suppress licking evoked by stimulation by generating a reach, and that this ability was influenced by VLS stimulation frequency.

### VLS stimulation extends drinking action

We also activated the VLS with a 500ms delay after beam break to stimulate during Drinking (**Figure 7A**). The result was sustained licking of the paw by mice for the duration of VLS stimulation (**Figure 7B & Video S5**). Increasing stimulation frequencies lead to increases in licking rate and a greater number of licks produced during Drinking (**Figure 7C & D**).

**Figure 7.**
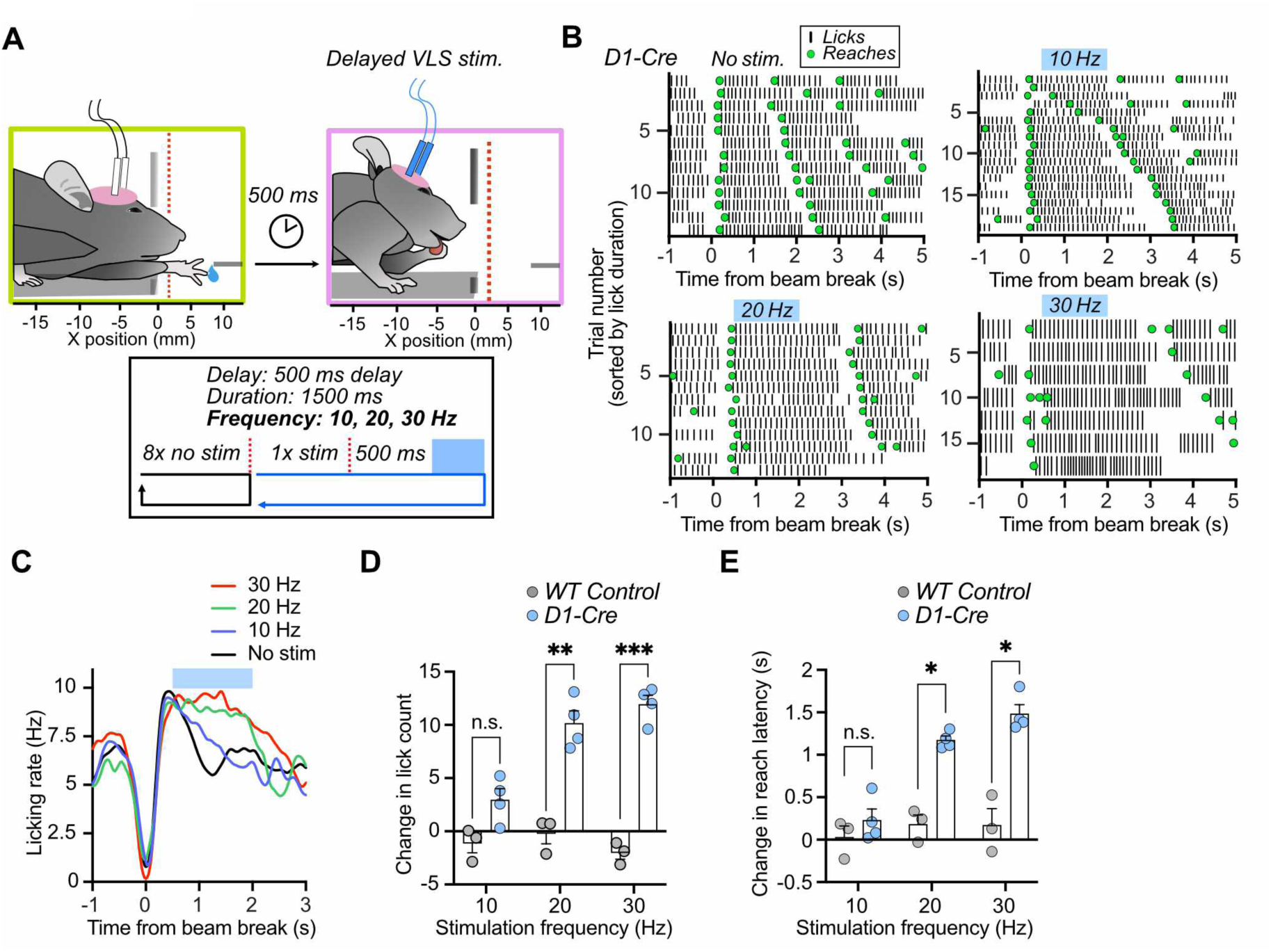
VLS direct pathway stimulation during Drinking extends drinking duration and delays reaching. **A)** Design. Laser stimulation lasting 1500 ms was delayed by 500 ms after the beam break, activating VLS D1 SPNs after reaching during Drinking. **B**) Example raster plots of licking aligned to the beam break during increasing stimulation frequency. Stimulation for 1500 ms extends licking and delays the next reach. **C**) Licking is extended by VLS D1 stimulation. Lines represent the mean licking rate for each stimulation condition (n = 4 mice per trace). **D**) Average change in lick number between stimulation onset and the onset of the next reach. (Two-way RM ANOVA: Significant main effects of Group (F(1,5) = 296.6, p < 0.0001) and Frequency (F(1.25,6.25**) =** 8.8, p = 0.02), Significant interaction (F(2,10) = 10.1, p = 0.004). Post-hoc tests found significant increases in lick number during 20 Hz (p = 0.003) and 30 Hz (p = 0.0001) trials. **E**) Higher stimulation frequencies delayed the next reach following stimulation. (Two-way RM ANOVA: Significant main effects of Group (F(1,5) = 39.8, p = 0.0015) and Frequency (F(1.85,9.26**) =** 28.3, p = 0.0001), Significant interaction (F(2,10) = 17.2, p = 0.0006). Post-hoc tests found significant increases in lick number during 20 Hz (p = 0.014) and 30 Hz (p = 0.023) trials.

Stimulation during Drinking delayed the initiation of the next reach with higher stimulation frequencies resulting in longer delays in mice generating the next reach (**Figure 7E**). Thus maintaining activity in the Drinking module was sufficient to delay reaching, suggesting that the Drinking module must be deactivated to allow other actions to progress. Together, our optogenetic stimulation results further support the evidence that a bidirectional lateral inhibitory mechanism resides in the basal ganglia.

## Discussion

Using the water-reaching task in mice, we uncovered a modular organization within the striatum that governs the precise temporal ordering of actions. Careful observation of the mouse performing reaching allowed us to characterize the rapid transitions between distinct effector groups during behavior. Using machine learning, we were able to parse this behavioral sequence precisely, allowing us to study the neural substrates corresponding to each component and how they are orchestrated in time ^25,26^.

### Modular Organization and Topographic Specificity

Our results demonstrate that the striatum contains distinct populations of striatal projection neurons (SPNs) that activate selectively during specific action components—aiming, reaching, and drinking—in a water-reaching task. These populations align with the known somatotopic organization of the corticostriatal projections, with DLS neurons active during Aiming and Reaching and VLS neurons engaged during Drinking. This topographic specificity supports previous findings that the DLS contribute to forelimb and trunk movements, while ventrolateral areas are more involved in orofacial movements^6,11^.

Complex behaviors can be broken down into discrete subgoals, each achieved by sending a velocity command associated with specific neural populations. The Aiming module is activated first, and deactivated as the head approaches the target, reflected in reduced head velocity. Because of the barrier, approaching with the head is not sufficient to obtain the water, and the remaining error in distance to water switches on the Reaching system, quantified by the transient increase in paw velocity. Like the Aiming population, once the target is acquired the Reaching module is suppressed as the Drinking modules is activated, as indicated by increased licking.

### Limitations in prior work on action sequencing

Prior work on behavioral sequencing in rodents use combinations of lever pressing to train arbitrary sequencing behaviors, such as left-left, right-right lever presses^17,18^. The experimenter-defined sequence can take several weeks to learn. Actions are recorded in the form of timestamps without monitoring the movement kinematics, and neural activity is described as a categorical feature (start/stop) due to the categorical nature of behavioral descriptions (press start/press end). It cannot show how neural activity continuously varies with the kinematic transitions to perform the task. These types of approaches have also resulted in ambiguity in interpreting optogenetic effects. For example, Tecuapetla et al. observed that direct pathway stimulation can delay action initiation, which we also observed (**Figure 7**). But the reason for this delay is unclear. It could have occurred because the stimulation promoted some other action that suppressed lever press initiation. Other studies have performed continuous monitoring of movement kinematics in arbitrarily trained skilled behavior^19^. The behavior sequences studied are not broken down and the subgoals are not defined. Consequently, neural activity cannot be related to the many possible action transitions used by the animals as they switch from one subgoal to the next in the sequence. Although it was concluded that the striatum represents kinematic variables, the way these relationships are used to generate specific sequences is not addressed. In our study, by continuously monitoring the kinematic transitions during a simple, easily understood sequence that is naturally produced by animals after limited training, we combined both types of approaches. This allowed us to reveal a process by which modular striatal activity associated with transitions between actions is selectively activated and suppressed to express these transitions in a particular order.

### Dynamic Interactions in Sequence Variations

The close alignment between SPN activity and kinematic profiles provides further support for striatum’s role in specifying detailed action kinematics in skilled movements^13,27–30^. SPNs showed rectified tuning, correlating with head and paw velocity toward the target but not during retraction. Drinking related SPNs, on the other hand, exhibited an antiphase relationship with paw velocity, peaking when paw movement was minimal during water consumption. This tight coupling between neural firing and effector kinematics indicates that striatal modules not only initiate actions but also modulate their execution, potentially optimizing movement efficiency or reward acquisition.

Analysis of double-reach trials revealed how the same striatal modules are flexibly recruited to facilitate trial-to-trial variations in behavioral sequences (**Figure 4**). During Interleaved double reaches, brief Drinking events interrupted Reaching. D-SPNs were activated as animals engaged in transiently licking the paw. Because of the suppression of R-SPNs during the activation of the Drinking module, R-SPNs must be activated a second time to generate the subsequent reach. In contrast, during rapid double reaches, R-SPN activity prior to the first reach was sufficient to sustain multiple reaches, since the module was not activated for the second reach. We assume that the downstream regions receiving striatal output are not discharged in the absence of any Drinking module activation. They can continue generating multiple movements, but these effects are not detected at the level of the striatum. The sustained reaching behavior may also prevent activation of the D-SPN activation and subsequently a delay in Drinking initiation. This pattern may be an example of “chunking” in the reaching task^1^.

These findings suggest that the basal ganglia’s modular organization is not rigidly recruited in a prescribed order. Instead, these systems are flexibly engaged to achieve the specific subgoals of each component of the sequence. Modular activation occurs if the competing system has previously discharged it but is not directly linked to the generation of every movement.

These systems are thus dynamically adjusted to the contextual demands to maintain sequence integrity.

### Reciprocal inhibition for pattern generation

The reciprocal inhibition mechanism identified here offers a mechanistic explanation for how the basal ganglia orchestrate complex behaviors. By suppressing competing actions, this mechanism ensures that each subgoal is executed in the correct order. We were able to discover a dynamic antagonism between different striatal modules, mediated by a reciprocal inhibition mechanism. During the reaching sequence, the activation of one module suppresses the activity of others, ensuring a strict serial order: aim, reach, drink. This was evident in the antiphase relationship between DLS and VLS activity, where DLS firing peaked during Reaching while VLS activity was suppressed, and vice versa during Drinking. Such reciprocal inhibition, possibly facilitated by lateral inhibition within the basal ganglia ensures that only one action dominates at a given time, preventing interference and maintaining the temporal precision of the sequence.

This account is supported by our optogenetic experiments. Activation of the VLS direct pathway provided causal evidence for the role of the Drinking module. Stimulation during Aiming triggered licking directed toward the water spout, suppressing head and paw movements associated with modules in the DLS (**Figure 5**). Furthermore, stimulation during Drinking maintained the active state of the Drinking module and prevented the expression of reaching (**Figure 7**) This suggests that the VLS drinking module can override other action modules, and must be discharged to facilitate other transitions, reinforcing the idea of reciprocal inhibition.

Conversely, in “Reaching Wins” and “Recovery” trials, reaching suppressed stimulation-evoked licking, indicating bidirectional inhibitory interactions (**Figure 6**). The frequency-dependent nature of these effects—higher stimulation frequencies increased licking and delayed reaching— further highlights the dynamic tuning of module interactions, with stronger activation amplifying one action while suppressing others.

While our findings provide strong evidence for modular organization and reciprocal inhibition, the precise circuitry underlying lateral inhibition requires further investigation. Both the striatum and the substantia nigra pars reticulata (SNr) have a lateral inhibition organization^31–33^. Their axon collaterals are recurrently connected and can produce inhibitory effects on each other. Within the striatum, lateral inhibition between SPNs is well established, but as these connections are characterized by low release probability, it remain unclear whether such inhibition is strong enough to suppress the firing of other SPNs^34,35^. Moreover, the axonal arbor of SPNs may not be large enough to affect SPNs that are far away, making it unlikely to have effective lateral inhibition if the modules are far apart. Because the Aiming and Reaching SPNs are found in close proximity to each other, it is possible that lateral inhibition between these systems can occur in the striatal microcircuit. Our results from activating the VLS revealed a robust suppression of head and paw movements, but the distances separating the large populations of Drinking neurons and the Aiming and Reaching neurons (**Figure 2**) make the collateral synapses between SPNs a less likely candidate for reciprocal inhibition.

Another possible mechanism is through lateral inhibition in the SNr^32^. Direct pathway activation is expected to inhibit many SNr output neurons, and disinhibit other SNr neurons via collateral inhibitory projections. Both increases and decreases in SNr activity were observed during direct pathway activation^36^. It is possible that the suppression of specific SNr neurons could increase the activity of others due to lateral inhibition.

In conclusion, our study provides evidence of spatially segregated functional modules that interact dynamically in real time to shape behavior. Our results reveal a modular, reciprocally inhibitory organization within the striatum, and shed light on how the brain orchestrates action sequences. The integration of high-resolution behavioral tracking with neural manipulations provides a new and highly efficient model for future research into the neural basis of motor control and sequence learning.

## STAR Methods

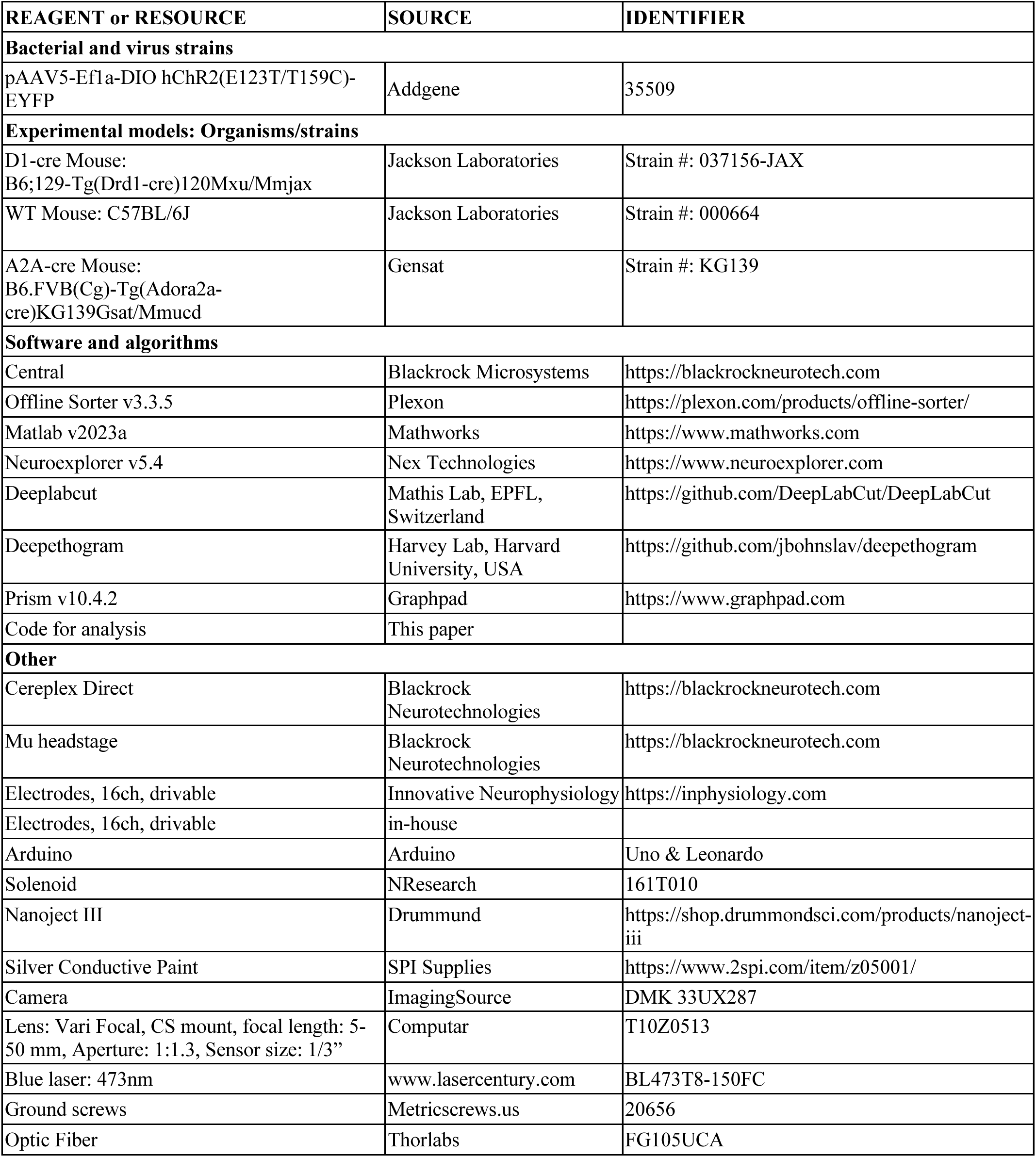
Key Resources Table.

## Resource availability

### Lead contact

Requests for additional information or resources can be directed to the lead contact, Henry Yin.

## Materials availability

The study did not generate any new or unique reagents.

## Data and code availability

All data reported in the study will be shared upon request from the lead contact. All original code will be uploaded to Duke’s Digital Repository.

## Experimental Model and Subject Details Animals

All animal procedures were approved by the Duke University Institutional Animal Care and Use Committee. Adult mice on the C57Bl6J background were used for all experiments. The study used 16 D1-cre mice (8 for optogenetics, 2 female, 6 male; 8 for electrophysiological recordings, 6 female, 2 male), 4 A2A-cre mice (for electrophysiological recordings, 2 female, 2 male), and 4 WT mice (for optogenetic controls, 4 male). Surgery was performed between 10 and 34 weeks after birth. Animals of the same sex and from the same litter were group housed on a 12:12hr light cycle.

## Method Details

### Adeno-associated virus

Optogenetic stimulation and control mice were injected with AAV5-Ef1a-DIO hChR2(E123T/T159C)-EYFP, Addgene catalog number 35509.

### Electrode and optical fiber surgery

Mice were anesthetized using 3% isoflurane in oxygen flowing at 0.6l/min and installed in a stereotaxic surgical frame on top of a thermal pad. Anesthesia was maintained between 1.5% and 0.5% over the course of surgery to maintain breathing rate and the absence of toe-pinch reflex. Eyes were kept lubricated with artificial tears. Hair was trimmed close to the scalp with scissors, and the skin was sterilized using iodine and ethanol wipes. Surgical tools were sterilized with a bead sterilizer for 30 seconds and allowed to cool on ethanol-soaked paper towels prior to beginning the incision. 4 sterile nickle ground screws (metricscrews.us, 20656) were implanted in the skull. For tagging, AAV2-ReaChR-Citrine was injected using a Nanoject III (Drummund) at the following coordinates: DLS: AP 0 mm, ML: +2.7 mm, DV: –4.0 & –3.25 VLS: AP 0 mm, ML: +2.7 mm, DV: –4.75 & –4.0. Injection volume: 200nL per site. The injector was removed and the ground wire for the 16 channel electrode arrays were wrapped around all ground screws. The connection was secured using conductive silver paint (SPI Supplies). Electrodes were lowered through a craniotomy made above the target location to the target depth. Implant coordinates for DLS were AP: 0 mm, ML: +2.7 mm, DV: –2.75 mm. Coordinates for VLS were AP: 0 mm, ML: +2.7 mm, DV: –3.75 mm.

Electrodes were affixed to the skull and ground screws with dental cement and a titanium headbar was attached to the cement cap for ease of headstage connection prior to recording. Mice were monitored for recovery for one week before beginning water deprivation and training. For optogenetic manipulations, surgical procedures were identical except that bilateral craniotomies were made above the sensorimotor striatum. AAV-ChR2 virus was infused bilaterally to the VLS prior to optic fiber implantation. Injection coordinates were AP: 0 mm, ML +/− 2.7 mm, DV: – 4.25 & –4.0 mm, with 150 nl of virus infused at each depth, beginning with the more ventral site, totaling 300 nl per hemisphere. After the final infusion, the injector was removed and bilateral optic fiber implants (100 um diameter optic fiber, 0.22 NA, Thorlabs) were lowered to a final DV position of –4.0 mm. Optic fiber implants were fixed to the skull and screws with dental cement. Mice were monitored for 1 week to ensure recovery from surgery and training began 2.5 weeks after surgery to allow for sufficient viral expression.

### Behavioral Setup and Video Recording

The arena was a plexiglass cube with a 1 cm wide opening on one side of the chamber. The opening extended from the floor of the arena to 4 cm above the floor. One corner of the arena was cut out to accommodate a camera lens that could be positioned perpendicularly to the main reaching axis, allowing accurate detection of movement initiations toward and away from the spout. A high-speed camera (ImagingSource) recorded behavior for the duration of the experimental session at 100 Hz. The camera’s GPIO pins were programmed to send a timestamp for each frame captured to a Blackrock Cereplex Direct, Blackrock Microsystems, Utah, USA). The spout was attached to a linear servo motor that allowed for precise changes in the spout’s distance from the arena. An infrared beam break was positioned across the opening to detect reaches. 10% sucrose in de-ionized water was delivered using a solenoid valve (NResearch) that regulated gravity assisted flow of water from a reservoir suspended above the arena. Food-grade polyethylene tubing (Cole Parmer) connected the reservoir to a blunted 21-guage syringe tip which served as the spout. Beam breaks, laser pulse times, rewards, and frame timestamps were logged together with electrophysiological recordings on the Blackrock Cereplex Direct.

### Task program

Arduino software ran the task and was programmed to deliver reward 500ms after the exit by the mouse from the beam break after each reach. Reward size was 10ul, which was controlled by adjusting the value opening duration and the height of the water reservoir. The program also counted the number of beam breaks in a row, and this counter was used to deliver optical stimulation on every 10 trials with a variable delay after bream break onset (see section on optogenetic stimulation below).

### Behavioral training

At least 1 week after surgery, mice were water deprived to approximately 90% baseline bodyweight. Animals were placed into the arena and the spout was positioned close to the aperture to allow the mice to lick the water. The spout was gradually moved further away to encourage reaching. Once animals performed the first reach, the spout was kept at a distance not reachable with the tongue. Mice could perform reaches on the first day. We encouraged the use of the left paw by positioning the spout slightly to the mouse’s right of the opening. This made the manual labeling of licking easier. Mice were trained daily for 30 minutes, during which time they could produce over 300 reaches. Mice were considered trained when they produced sustained bouts of reach behavior. Mice produced sustained bouts of repeated Aim, Reach, Drink cycles at the latest by day 5.

### Kinematics

A DeepLabCut model was trained to identify the position of the nose and the left paw in each frame in x and y coordinates. Coordinates were converted from pixels into millimeter values by applying a conversion factor based on a known straight edge in the video. Velocity was calculated by taking the difference between the position value of a given frame and the position value of the prior frame and dividing this result by the video’s inter-frame interval. Analysis was only performed on the velocity of the x position – positive velocity values reflected movements out of the arena toward the spout and negative values reflect movements into the box, away from the spout.

### Action assignments

a DeepEthogram model was trained to assign an action to each frame in the video. The model was manually trained to label frames as reflecting Aim when mice ceased licking from the hand and began moving their nose toward the spout. The model was trained to assign frames as Reach when the hand ceased shaking and would begin a smooth trajectory toward the spout and to assign frames to the Drinking action when the mouse began licking the paw after grabbing the water. Licking was trained when frames showed extension of the tongue while the mouse was in the Aim posture. Scavenge was assigned when mice licked the floor of the arena. The Out label was assigned to frames that showed any other type of movement that could not be assigned to any other category or when the mouse was not in the frame.

### Licking tracking

Perievent video clips were generated around relevant experimental events and were then imported into a manual frame labeling software (Boris). Licks were scored as the point of maximum tongue extension.

### In-vivo recording and spike sorting

Neural activity was recorded from freely behaving animals using a Blackrock Cereplex Direct (Blackrock Neurotech, Utah, USA) and a 16 channel Cereplex mu headstage. Mice were connected to the headstage and placed into the reaching chamber without the possibility of obtaining reward while units were sorted online using Central (Blackrock Neurotech, Utah, USA). After sorting, the reward solenoid was activated, and video recording began once the mice began reaching. Electrodes were advanced by 50 um after each recording session. Offline Sorter (Plexon) was used after the recording to confirm that units sorted on-line were well isolated. Recording tracts were constructed by taking the depth of the first and last recording session. Final electrode positions were confirmed post-mortem.

### Optogenetic stimulation

473 nm wavelength laser was coupled to a rotating splitting commutator (Doric) which was then connected to two connecting optic fibers. Power settings were set to 5 mW on each connecting fiber before each experiment. Mice were connected bilaterally to the connecting fibers via their implants. The fibers were covered with plastic tubing to minimize flashing visual stimuli and the mice placed into the reaching chamber. An infrared beam break (Adafruit) was positioned so that the beam crossed just in front of the reaching aperture. The beam break was further adjusted for each animal so that the movement of the nose through the aperture would break the beam. Beam breaks were counted using an Arduino Leonardo, which sent a trigger signal to the laser upon the 9^th^ beam break crossing. Laser pulse duration was 10ms. Animals could receive stimulations at different frequencies (10hz, 20hz, or 30hz) in each experimental session if their motivation allowed for continued attempts to reach.

We aimed to deliver at minimum 10 stimulation trials per condition. For experiments with delayed stimulation, animals received the same stimulation parameters but after a 500ms delay after the beam break to stimulate during Drinking. WT control animals received all the same treatments. All analyses were performed relative to the beam break.

## Quantification and Statistical Analysis

### Spiny projection neuron classification

All spike waveforms for each unit were sampled for 1.6ms at 30kHz. For each unit, the waveforms from the entire session were averaged to obtain the unit’s average waveform. Waveforms from all units in the dataset were arranged into a single matrix with rows corresponding to neurons and columns corresponding to time. Waveforms were normalized using z-score function in Matlab to center the waveform amplitude around zero and to scale this amplitude to have a standard deviation of 1. Principal component analysis was applied to the normalized waveform matrix. The top three principal components were selected based on having the highest eigenvalues. Waveforms were projected onto these components to obtain a three-dimensional representation of each waveform in PCA space. Waveforms were clustered based on these projections using hierarchical clustering with Ward’s linkage method and 2 expected clusters. The mean firing rate and full-width-half-max (FWHM) was calculated for each unit.

### Cell population functional classification

Mean firing rates for each SPN aligned to the onset of Reach DEG labels. Reach onset times included in the analysis had to meet the following criteria: they had to be preceded by Drinking and Aiming and followed by Drinking. This was done to analyze reaches that occurred within bouts and excluded departures and entries into the task.

Neurons were filtered based on their classification as SPNs. Firing rates were binned at 50ms intervals and smoothed using a gaussian kernel (12 bins). For DLS the analysis window was –1 to +1 s around reach onset. For VLS, it was –1 to +1.5 s to incorporate drink related activity.

Neurons with spiking activity on less than 10% of trials were excluded from analysis as non-modulated. To identify functional groups of neurons, firing rate profiles in the analysis window were normalized using z-score standardization. Hierarchical clustering with Ward’s linkage was applied to the normalized data, using a Euclidean distance metric and a predefined number of clusters (k = 5).

### Firing rate analysis

To detect reductions and increases in activity, we computed the zscore. Neural activity baselines were first calculated by taking the mean firing rate for a window from – –30 to –2 seconds prior to all reach onsets in the session. The baseline standard deviation was derived from the distribution of these firing rate estimates across trials. To calculate zscore, firing rates aligned to action onset were baseline subtracted and then divided by the baseline firing rate standard deviation. Activity was suppressed if the zscore fell below 0 and was increasing of activity was above 0. To compare changes in firing rates for Aiming, Reaching, and Drinking SPNs, we aligned the activity of the cells to each of the 3 actions in the sequence. For Aiming and Reaching SPNs, we calculated the average zscore for the 100ms period prior to Aim and Reach onset to quantify their activity during these actions. To quantify their firing activity related to Drinking, we took the average zscore for the first 500ms after Drinking. For Drinking SPNs, we calculated the average zscore of their activity for the 500ms after Aiming, Reaching, and Drinking.

### Tuning for kinematics

For each session, we generated peri-event matrices for the velocity of the head and paw markers, aligned to all reach actions in the session, using a window of –1 to 1 and 25 ms bins. For each unit in the session, we generated peri-event firing rate matrices aligned to the onset time of all reach actions in the session, also using a window of –1 to 1 seconds and 25 ms time bins. The behavior matrices were linearized to produce 2 vectors of head and paw velocity from all reaches concatenated end to end. The same linearization was performed for the peri-event firing rate matrix for each unit to produce a firing rate vector. A cross correlation was performed for each unit to test for the optimal alignment of firing rate with each kinematic variable. The tuning curve was constructed by aligning the firing rate vector to the kinematic variable that had the lowest positive lag behind the unit activity and shifting the firing rate vector according to that lag. Velocity values were then binned into 18 bins from most negative to most positive, and the firing rate for all observations of velocity at each binned value were averaged together to produce the tuning curve for each unit. The resulting single unit tuning curve was then scaled from 0 to 1 and all units of a given class were averaged together to generate the population tuning curves. No cross-correlation lag adjustments were applied for Drink-SPNs.

### Double reaches

Although Deepethogram could detect reaching actions, it sometimes fails to detect individual reaches within a bout of multiple reaches. To enhance the resolution of reach event detection, we identified individual reaches by finding peaks of paw position when they exceeded 4 mm past the reaching port. We first computed the inter-reach-interval (IRI) between successive reach events. Trials with an IRI of < 750 ms were classified as candidate double reaches. Trials were divided into Rapid or Interleaved categories based the IRI between the reaches in the double-reach event. Interleaved type trials had IRIs between 250 and 750ms whereas trials with reaches having an IRI less than 250ms were classified as rapid reach trials. Sessions had to contain both types of trials and had to contain at least 3 Rapid Reach trials for kinematics, Deepethogram labels, and firing rates to be further analyzed. Neural activity was aligned to the first and second reach peak events in a double-reach bout and binned with 25ms windows. The zscore was computed as described above. Reductions of zscore below 0 were considered activity suppression and increases above 0 were considered activation.

### Statistical analysis

All analyses were performed using Neuroexplorer, Matlab, and GraphPad Prism. Sample sizes (number of mice or number of units) and statistical tests used are included in the figure legends. The significance level of the p-value was set at p < 0.05. Data are reported as +/− SEM. No steps were taken to confirm assumptions of the statistical tests used.

**Figure S1.**
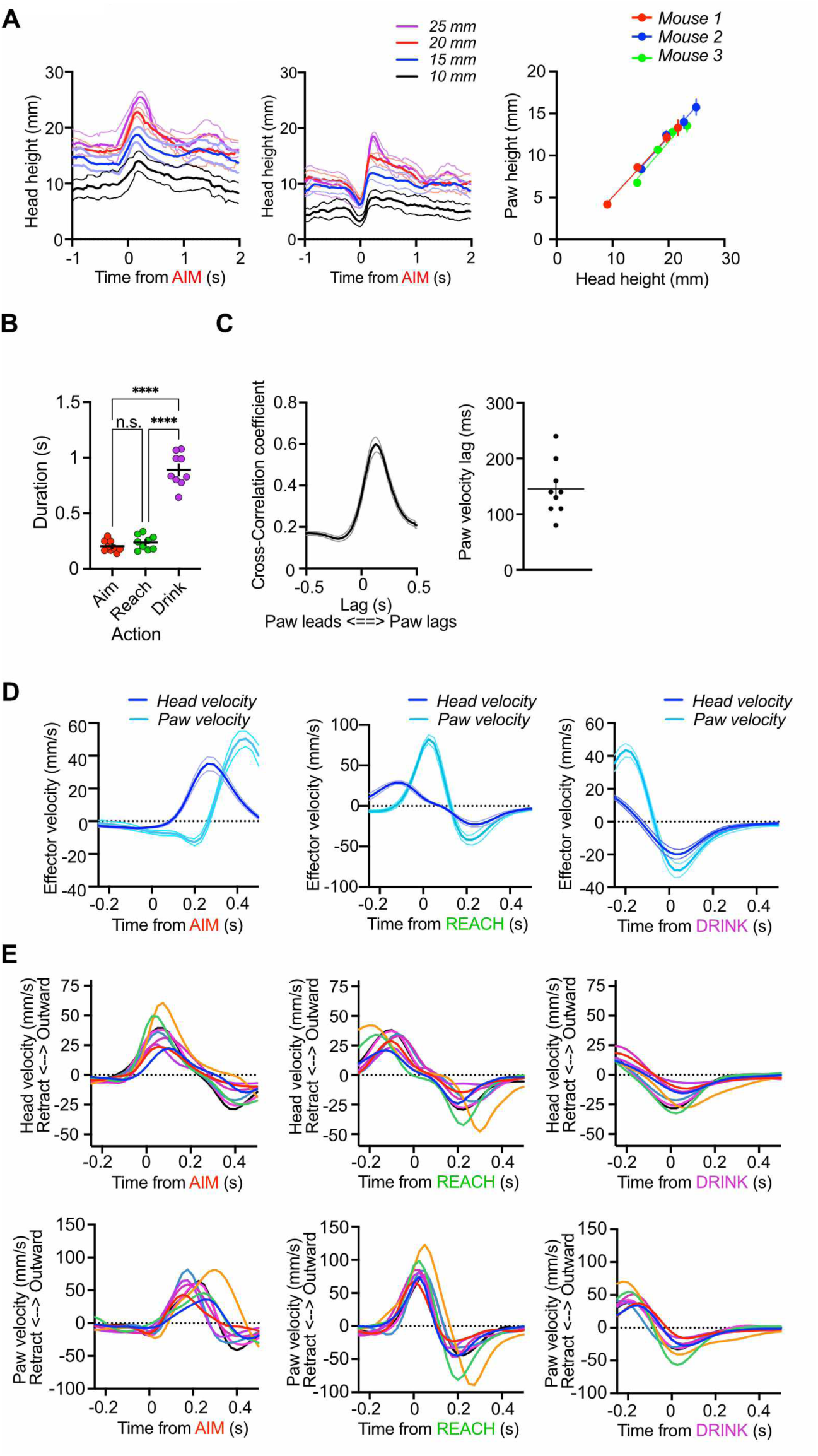
Paw velocity lags head velocity. **A)** Head (*Left*) and paw (*Middle*) height as a function of changing spout height above the base of the reaching chamber. *Right*: Head and paw height are covaried to access the water spout (n = 3). **B)** Reaching occurred approximately every 1.25 seconds (n = 9). One-way RM ANOVA (F(1.4, 11.2) = 122.9, p < 0.0001. Post-hoc tests showed that Aiming and Reaching did not differ in duration (p = 0.48), but were both shorter than Drinking (p < 0.0001). **C**) *Left*: Mean cross correlogram showing that paw velocity consistently lags head velocity *Right*: Mean lag was 145.6 ms. **D**) *Left*: Average velocity of the head and paw aligned to Aiming onset (n = 9 mice). Increasing head velocity coincides with low paw velocity. *Middle*: Average velocity of the head and paw aligned to Reaching onset (n = 9 mice). Increasing paw velocity coincides with low head velocity. *Right*: Average velocity of the head and paw aligned to Drinking onset (n = 9). **E**) Mean velocity traces of head (*Top row*) and paw (*Bottom row*) for 9 representative animals aligned to Aim, Reach, and Drink onsets. Data is the same data that is shown in averaged form in *E*. Related to Figure 1.

**Figure S2.**
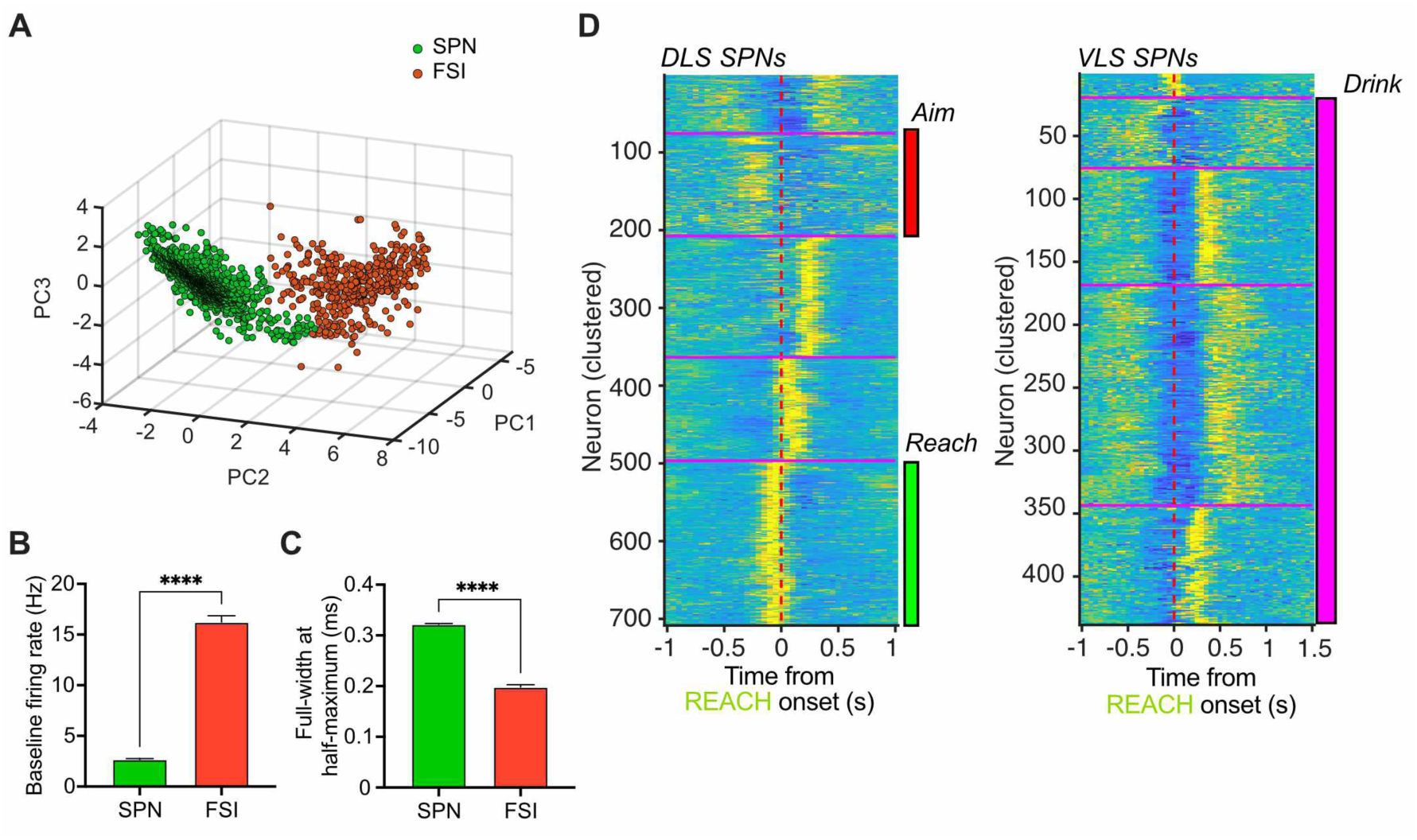
Cell classification. **A**) Projection of waveforms from striatal neurons into principal component space. Cells were separated into two groups, putative SPNs and FSIs. **B**) Average baseline firing rates for SPNs was lower than for FSI clusters. (unpaired t-test, p < 0.0001). **C**) Waveforms of SPNs were wider than for FSIs, as measured by the Full-width at Half-Max (unpaired t-test p < 0.0001). **D**) Heatmaps of population firing rates of SPNs recorded in the DLS and the VLS. Firing rate was aligned to Reach onset and, averaged across trials, Z-score normalized and clustered using agglomerative hierarchical clustering using a cluster number of 5. Task-relevant Aiming, Reaching, and Drinking related populations are highlighted with colored bars. Related to Figure 2.

**Figure S3.**
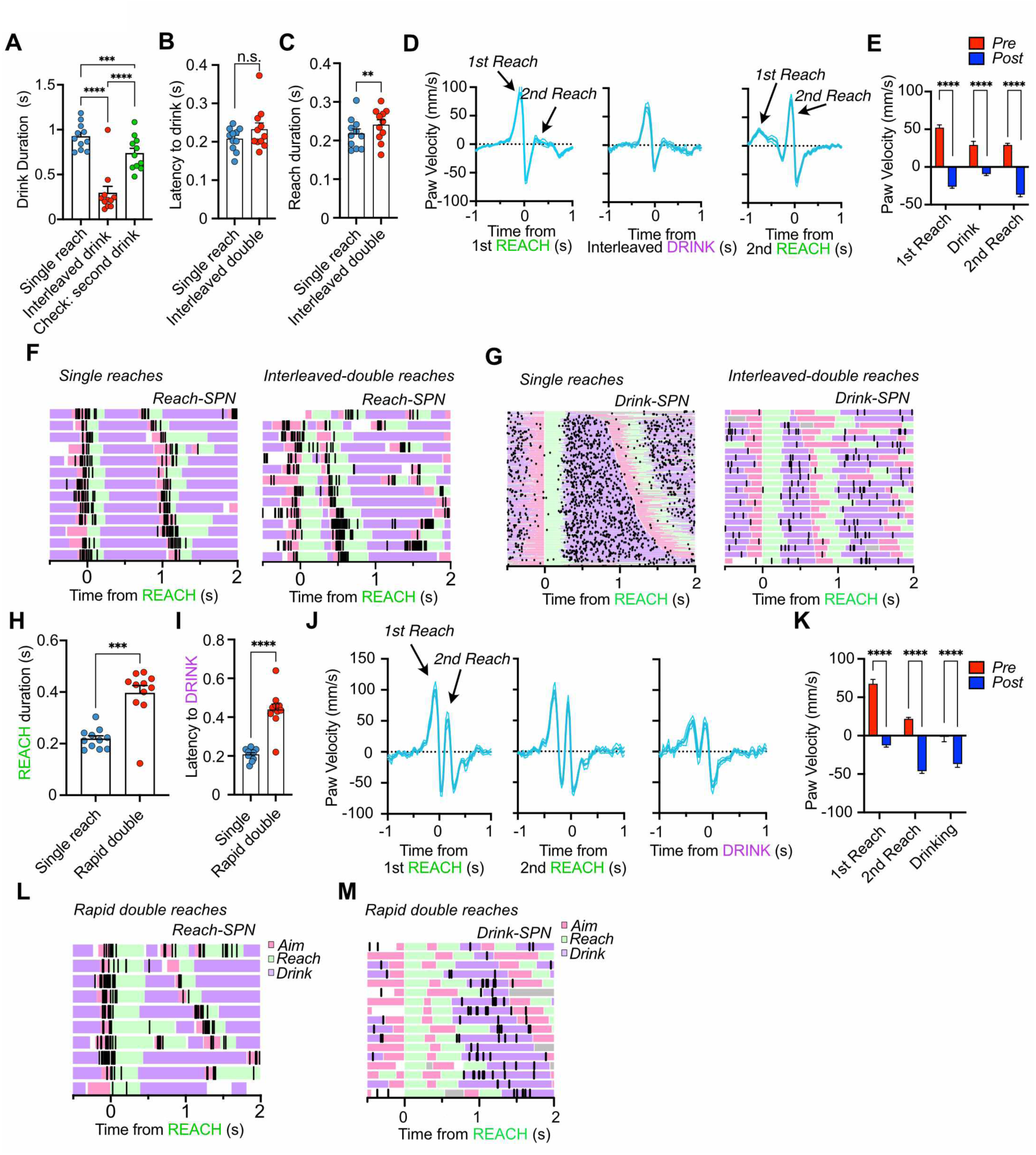
Details of Double-reach events. **A**) A significant one-way RM ANOVA found a significant effect of drink event on duration (F(1.526, 15.26) = 85.65, p < 0.0001). The duration of interleaved drinking was lower than drinking after single reaches (p < 0.0001) and lower than the second drink event during Interleaved-double reaches (p < 0.0001). The second drinking event during Interleaved-double trials was also shorter in duration than drinking during single reaches (p = 0.0006). **B**) Mice did not change the latency to initiate drinking during Interleaved-double trials. **C**) Reach duration was slightly longer during Interleaved-double trials. **D**) Average paw velocity aligned to the 1^st^ reach, interleaved drink, and 2^nd^ reach during Interleaved-double trials (n = 11mice). **E**) Average paw velocity aligned to each action in Interleaved-double trials. Paw velocity was averaged prior to each action (−250 to 0ms) and after each action (0 to 250ms). A two-way RM ANOVA found significant main effects of action (F(2,20) = 27.6, p < 0.0001) and analysis interval (F(1,10) = 188.3, p < 0.0001) and a significant interaction between Action and Trial Type (F(2,20) = 29.83, p < 0.0001). Post-hoc tests found reduced paw velocity after the 1^st^ reach (p < 0.0001), after interleaved drink (p < 0.0001) and after the 2^nd^ reach (p < 0.0001). **F**) Raster plots and Deepethograms for example R-SPNs during Single reaches (*Left*) and Interleaved double reaching (*Right*). **G**) Raster plots and Deepethograms for example D-SPNs during single reaches (*Left*) and Interleaved-double reaching (*Right*). **H**) The duration of reaching was increased during Rapid double reaching (p = 0.0001). **I**) Rapid double reaching delayed drink onset latency (paired t-test, p < 0.0001). **J**) Average paw velocity aligned to the 1^st^ reach, 2^nd^ reach, and drinking action during Rapid-double trials (n = 11mice). **K**) Average paw velocity aligned to each action in Rapid-Double trials. Paw velocity was averaged prior to each action (Pre: –150 to 0ms) and after each action (Post: 0 to 150ms, note narrower time window because of the speed of the movements). A two-way RM ANOVA found significant main effects of Action (F(2,20) = 94.75, p < 0.0001) and Analysis interval (F(1,10) = 114, p < 0.0001) and a significant interaction between Action and Trial Type (F(2,20) = 27.78, p < 0.0001). Post-hoc tests found reduced paw velocity after the 1^st^ reach (p < 0.0001), after the 2^nd^ reach (p < 0.0001) and after Drinking (p < 0.0001). **L**) Raster plot and Deepethogram for example R-SPN during Rapid-double reaching. **M**) Raster plot and Deepethogram for example D-SPN during Rapid-double reaching. Related to Figure 4.

**Figure S4.**
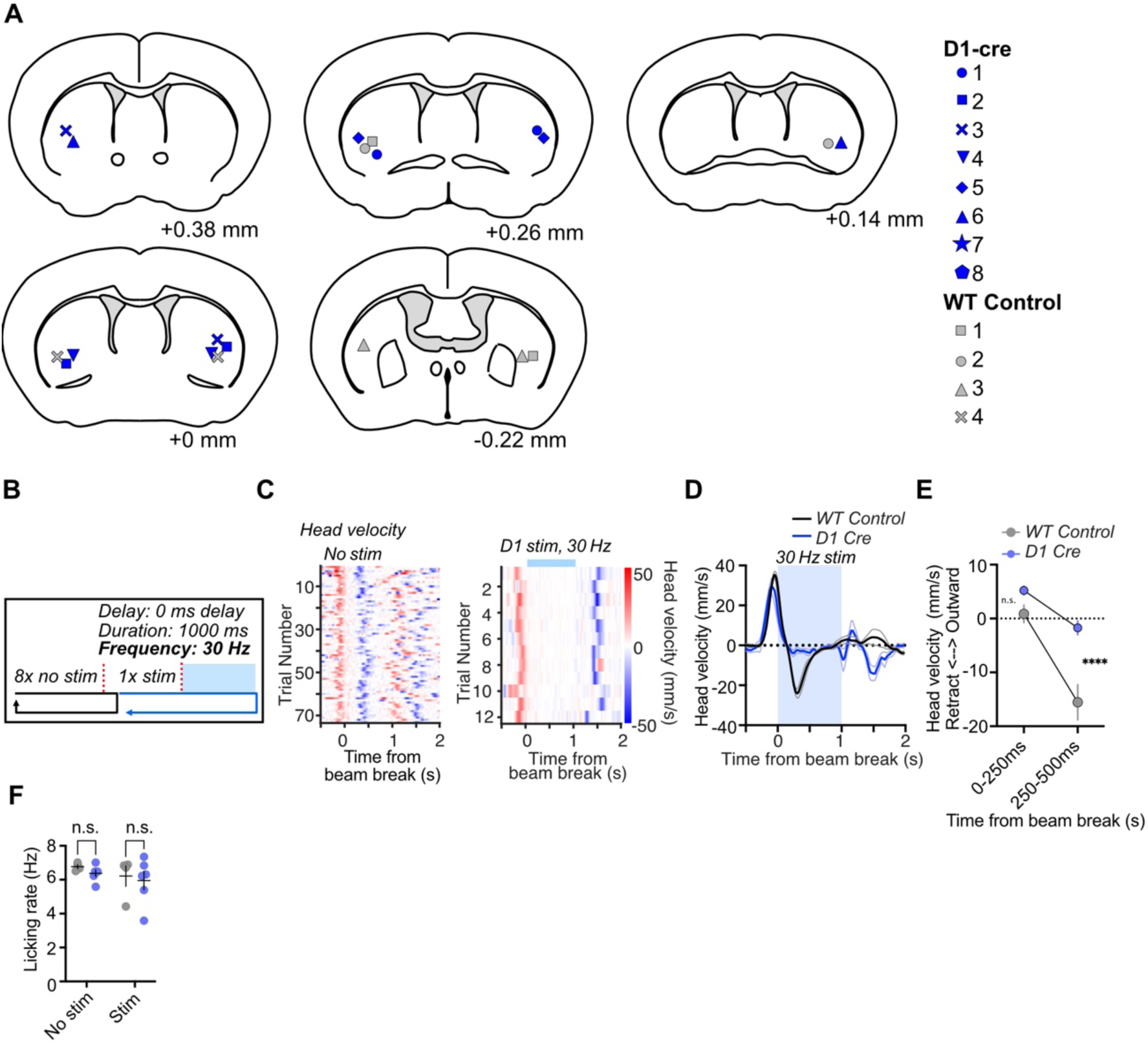
VLS direct pathway stimulation reduces head velocity. **A**) Locations of optical fiber tips across all D1-cre and WT control mice. **B**) Stimulation design. 30 Hz stimulation was activated at the time of beam break. **C)** Representative heatmaps showing the loss of negative velocity of the head during VLS direct pathway activation. **D**) Average head velocity time series of all subjects showing that D1pathway stimulation suppresses head movement (n = 6 D1-cre and 4 WT Control mice). **E**) VLS direct pathway stimulation prevents head retraction in D1-cre mice. A two-way ANOVA found significant main effects of Time Point (F(1,8) = 27.6, p = 0.0008) and Stimulation (F(1,8) = 67.7, p < 0.0001) on head velocity. **F**) A two-way ANOVA did not find significant effects of Stimulation (F(1,8) = 1.18, p = 0.3) or Group (F(1,8) = 0.64, p = 0.44) on lick rate. Related to Figure 5.

**Figure S5.**
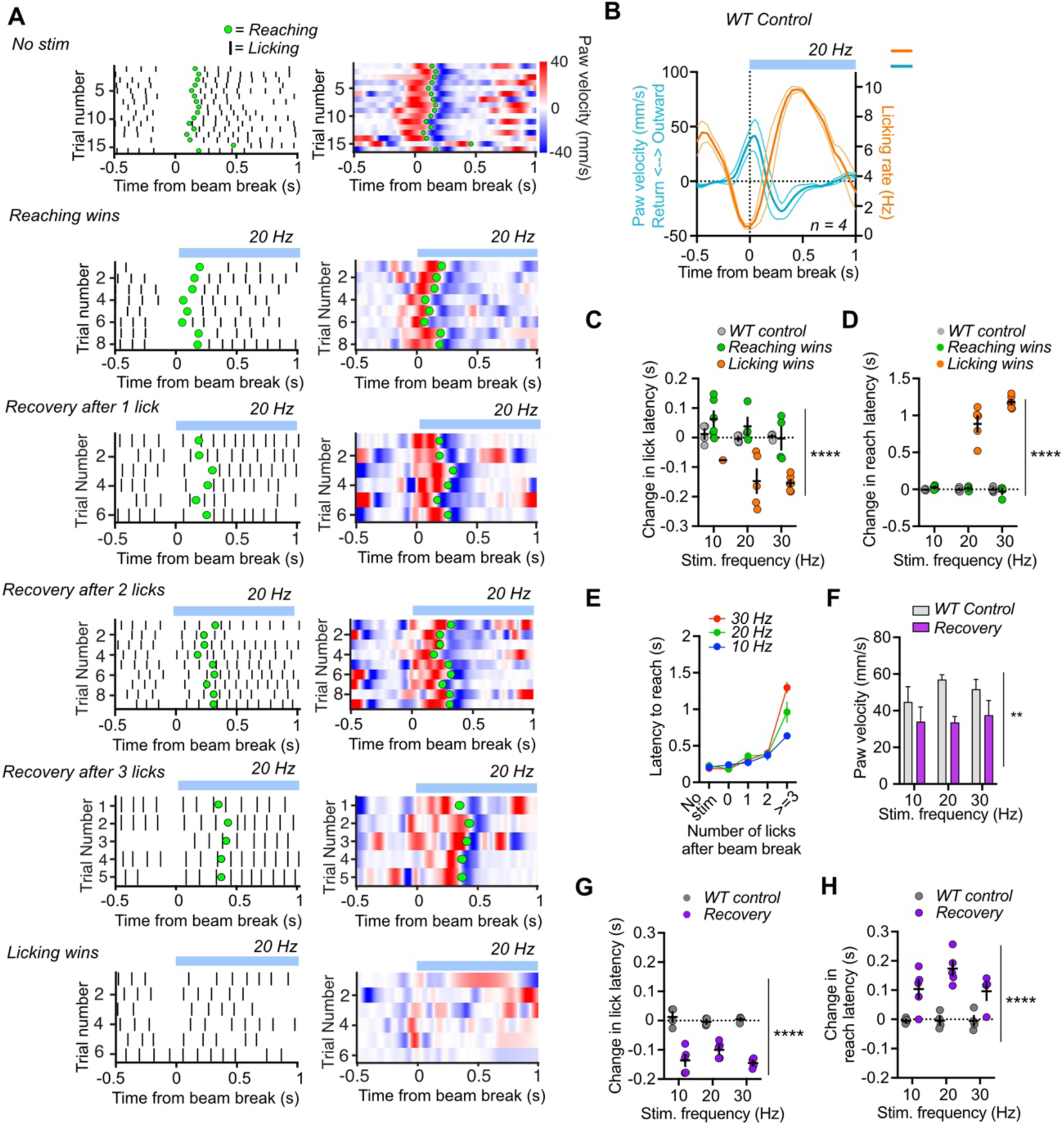
Additional details of the suppression of stimulation-evoked licking by reaching. **A**) Licking rasters and paw velocity heatmaps for a representative mouse receiving 20Hz stimulation. All trial types are shown, No stim, Reaching Wins, Recovery, and Licking Wins. Recovery trials are divided based on the number of licks produced prior to the Recovery reach event. Note slight increases in paw velocity during the initial licks. Mice are attempting to move the paw during stimulation. Reach events are denoted by the green dots. **B**) Average Paw velocity and Licking rate for control mice during 20 Hz stimulation (n = 4 mice). **C**) Licking latency was reduced during Licking wins trials (Two-way ANOVA, F(2,29) = 19.63, p < 0.0001). **D**) Reaching latency was increased during Licking wins trials (Two-way ANOVA, F(2,32) = 256, p < 0.0001). **E**) Reaching was delayed by increasing lick number. Two-way ANOVA, significant main effects of Lick number (F(4,52) = 64, p < 0.0001) and Frequency (F(2,52) = 3.7, p = 0.03), significant interaction (F(8,52) = 4.5, p = 0.0003). Post-hoc tests found that reaching was significantly delayed compared to no-stim trials if the mice generated 2 licks (p = 0.045) or more than 3 licks (p < 0.0001) before reaching was initiated. **F**) Paw velocity at the time of beam break was lower on Recovery trials where licking would temporarily dominate. Two-way ANOVA, significant main effect of Group (F(1,20) = 9.4, p = 0.0061). **G**) Licking latency is reduced during Recovery trials (Two-way ANOVA, main effect of Trial type (F(1,20) = 149, p < 0.0001). **H**) Reaching latency is increased during Recovery trials (Two-way ANOVA, main effect of Trial type (F(1,20) = 45, p < 0.0001). Related to Figure 6.

## Supplemental Videos

**Video S1: Motion tracking and action segmentation during reaching in mice**

**Video S2: Flexible alterations in sequences during double reach trials**

**Video S3: VLS direct pathway stimulation promotes licking and suppresses reaching**

**Video S4: Reaching during VLS direct pathway stimulation suppresses light-evoked licking**

**Video S5: VLS direct pathway stimulation prolongs Drinking and delays Reaching**

